# Functional genomics of cattle through integration of multi-omics data

**DOI:** 10.1101/2022.10.05.510963

**Authors:** Hamid Beiki, Brenda M. Murdoch, Carissa A. Park, Chandlar Kern, Denise Kontechy, Gabrielle Becker, Gonzalo Rincon, Honglin Jiang, Huaijun Zhou, Jacob Thorne, James E. Koltes, Jennifer J. Michal, Kimberly Davenport, Monique Rijnkels, Pablo J. Ross, Rui Hu, Sarah Corum, Stephanie McKay, Timothy P.L. Smith, Wansheng Liu, Wenzhi Ma, Xiaohui Zhang, Xiaoqing Xu, Xuelei Han, Zhihua Jiang, Zhi-Liang Hu, James M. Reecy

## Abstract

Functional annotation of the bovine genome was performed by characterizing the spectrum of RNA transcription using a multi-omics approach, combining long- and short-read transcript sequencing and orthogonal data to identify promoters and enhancers and to determine boundaries of open chromatin. A total number of 171,985 unique transcripts (50% protein-coding) representing 35,150 unique genes (64% protein-coding) were identified across tissues. Among them, 159,033 transcripts (92% of the total) were structurally validated by independent datasets such as PacBio Iso-seq, ONT-seq, *de novo* assembled transcripts from RNA-seq, or Ensembl and NCBI gene sets. In addition, all transcripts were supported by extensive independent data from different technologies such as WTTS-seq, RAMPAGE, ChIP-seq, and ATAC-seq. A large proportion of identified transcripts (69%) were novel, of which 87% were produced by known genes and 13% by novel genes. A median of two 5’ untranslated regions was detected per gene, an increase from Ensembl and NCBI annotations (single). Around 50% of protein-coding genes in each tissue were bifunctional and transcribed both coding and noncoding isoforms. Furthermore, we identified 3,744 genes that functioned as non-coding genes in fetal tissues, but as protein coding genes in adult tissues. Our new bovine genome annotation extended more than 11,000 known gene borders compared to Ensembl or NCBI annotations. The resulting bovine transcriptome was integrated with publicly available QTL data to study tissue-tissue interconnection involved in different traits and construct the first bovine trait similarity network. These validated results show significant improvement over current bovine genome annotations.

## Introduction

Domestic bovine (*Bos taurus*) provides a valuable source of nutrition and an important disease model for humans (Roth and Tuggle 2015). Furthermore, cattle have the greatest number of genotype associations and genetic correlations of the domesticated livestock species, which means they provide an excellent model to close the genotype-to-phenotype gap. Therefore, the accurate identification of the functional elements in the bovine genome is a fundamental requirement for high quality analysis of data informing both genome biology and clinical genomics.

Current annotations of farm animal genomes largely focus on the protein-coding regions and fall short of explaining the biology of many important traits that are controlled at the transcriptional level (Beiki et al. 2019). In humans, 88% of trait-associated single nucleotide polymorphisms (SNP) identified by genome-wide association studies (GWAS) are found in non-coding regions (Hindorff et al. 2009). Therefore, elucidating non-coding functional elements of the genome is essential for understanding the mechanisms that control complex biological processes.

Untranslated regions play critical roles in the regulation of mRNA stability, translation, and localization (Jereb et al. 2018), but these regions have been poorly annotated in farm animals (Schurch et al. 2014; Beiki et al. 2019). A recent study of the pig transcriptome using single-molecule long-read isoform sequencing technology resulted in the extension of more than 6000 known gene borders compared to Ensembl or National Center for Biotechnology Information (NCBI) annotations (Beiki et al. 2019).

Small non-coding RNAs, such as microRNAs (miRNA), are known to be involved in gene regulation through post-transcriptional regulation of expression via silencing, degradation, or sequestering to inhibit translation (Ambros 2004; Bartel 2004; Yates et al. 2013). The number of annotated miRNAs in the current bovine genome annotation (Ensembl release 2018-11; 951 miRNAs) is much lower than the number reported in the highly annotated human genome (Ensembl release 2021-03; 1,877 miRNAs).

This study applied a comprehensive set of transcriptome and chromatin state data from 47 cattle tissues and cell types to identify previously unannotated genes and improve the annotation of thousands of protein-coding and non-coding genes. Predicted novel genes and transcripts were highly supported by independent Pacific Biosciences single-molecule long-read isoform sequencing (PacBio Iso-Seq), Oxford Nanopore Technologies sequencing (ONT-seq), Illumina high-throughput RNA sequencing (RNA-seq), Whole Transcriptome Termini Site Sequencing (WTTS-seq), RNA Annotation and Mapping of Promoters for the Analysis of Gene Expression (RAMPAGE), chromatin immunoprecipitation sequencing (ChIP-seq), and Assay for Transposase-Accessible Chromatin using sequencing (ATAC-seq) data. The transcriptome data was integrated with publicly available Quantitative Trait Loci (QTL) and gene association data to construct the first bovine trait similarity network that recapitulates published genetic correlations. Thus, it may be possible to begin to examine the genetic mechanisms underlying genetic correlations.

## Results

The diversity of RNA and miRNA transcript diversity among 47 different bovine tissues and cell types was assessed using miRNA-seq and poly(A)-selected RNA-seq and miRNA-seq data. Most of the tissues studied were from Hereford cattle closely related to L1 Dominette 01449, the individual from which the bovine reference genome (ARS-UCD1.2) was sequenced. The 47 tissues and cell samples included follicular cells, myoblasts, five mammary gland samples from various stages of mammary gland development and lactation, eight fetal tissues (78-days of gestation), eight tissues from adult digestive tract, and 16 other adult organs. A total of approximately 4.1 trillion RNA-seq reads and 1.2 billion miRNA-seq reads were collected, with a minimum of 27.5 million RNA-seq and 9.3 million miRNA-seq reads from each tissue/cell type (average 87.8 ± 49.7 million and 27.6 ± 12.9 million, respectively) (Supplemental file 1: Fig. S1 and Supplemental file 2).

### Transcript level analyses

A total of 171,985 unique transcripts (76% spliced) were identified (Table 1) with a median of 51,231 transcripts per tissue. The median length of exons was 137 nt, and that of introns was 1,428 nt. Exonic length of transcripts (region of transcript covered by exons after collapsing transcript exons) was significantly longer (p-value < 2.2e-16) in spliced transcripts (median of 1,651 nt) compared to unspliced transcripts (median of 513 nt). There was a median of 9.1 exons per spliced transcript, and all of the predicted acceptor and donor splice sites conformed to the canonical consensus sequences. All of the predicted splice junctions across tissues were supported by RNA-seq reads that spanned the splice junction, substantiating the accuracy of the transcript definition from RNA-seq reads.

**Table 1.** Summary of detected transcripts/genes

| Feature | Annotation<sup>1</sup> |  |  |
| --- | --- | --- | --- |
|  | Cattle FAANG | Ensembl | NCBI |
|  |  | (Release 2021-03) | (Release 106) |
| Number of genes | 35,150 (21,193) | 27,607 (21,880) | 35,143 (21,355) |
| Number of transcripts | 171,985 (85,658) | 43,984 (37,538) | 83,195 (47,280) |
| Number of spliced transcripts | 130,531 | 37,299 | 73,423 |
| Number of transcripts per gene | 4.9 | 1.5 | 2.3 |
| Median number of 5' UTRs per gene | 2 | 1 | 1 |
| Median number of 3' UTRs per gene | 1 | 1 | 1 |

<sup>1</sup>Numbers in parentheses indicate the number of protein-coding genes/transcripts.

A total of 31,476 transcripts appeared tissue-specific by virtue of being assembled from RNA-seq reads in just a single tissue, but 20,100 of those transcripts (64%) were actually expressed in multiple tissues according to long-read Iso-seq data. Thus, reliance solely on assembled transcripts in a given tissue to predict a tissue transcript atlas may overestimate tissue specificity due to a high false-negative rate for transcript detection. To solve this problem of over-prediction of tissue specificity, we marked a transcript as “detected” in a given tissue only if (1) it had been assembled by RNA-seq data in that tissue; or (2) it had been detected by Iso-seq data in another tissue, but all splice junctions were validated using RNA-seq reads in the tissue of interest with an expression level more than 1 RPKM (see Methods section). This resulted in 15,562 apparently tissue-specific transcripts (9%) and 156,423 transcripts (91%) detected in more than one tissue (Fig. 1A), among which 9,125 transcripts (5%) were found in all 47 tissues examined.

**Figure 1.**
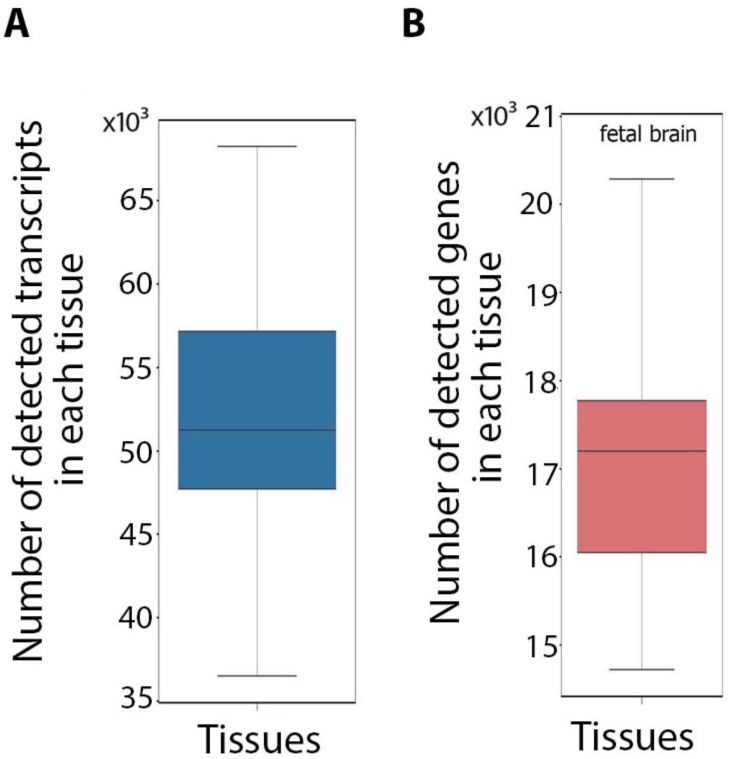
Distribution of the number of detected transcripts (A) and genes (B) across tissues.

The unique transcripts identified were equally distributed between 85,658 (50%) protein-coding transcripts and 86,327 (50%) non-coding transcripts (ncRNAs) (Fig. 2). Non-coding transcripts were further classified as long non-coding (lnc) RNAs (56%), nonsense-mediated decay (NMD) transcripts (38%), non-stop decay (NSD) transcripts (5%), and small nuclear (sn) RNAs (1%). While the majority of detected transcripts in each tissue were protein coding (median of 62% of tissue transcripts), NMD transcripts (median of 14.58% of tissue transcripts) and antisense lncRNAs (median of 12% of tissue transcripts) each made up more than 10% of the transcripts (Supplemental file 1: Fig. S2A and B, Supplemental file 3 and 4). Fetal muscle and fetal gonad tissues showed the highest proportion of antisense lncRNAs compared to that observed in other tissues (Supplemental file 1: Fig. S2B) and around 60% of antisense lncRNAs (17,982 transcripts) were detected from these two tissues. Compared to non-coding transcripts, protein-coding transcripts were more likely to have spliced exons (p-value < 2.2e-16) and were detected in a higher number of tissues (median of 11 tissues for protein-coding transcripts versus six tissues for non-coding transcripts; p-value < 2.2e-16) (Additional file1: Fig. S2C). The lncRNAs had a significantly lower splice rate (36%) compared to other non-coding transcripts (p-value < 2.2e-16). Splice rate was highest (70%) in sncRNAs (p-value < 2.2e-16; NMD transcripts were not included in this analysis, as they were all spliced transcripts by definition).

**Figure 2.**
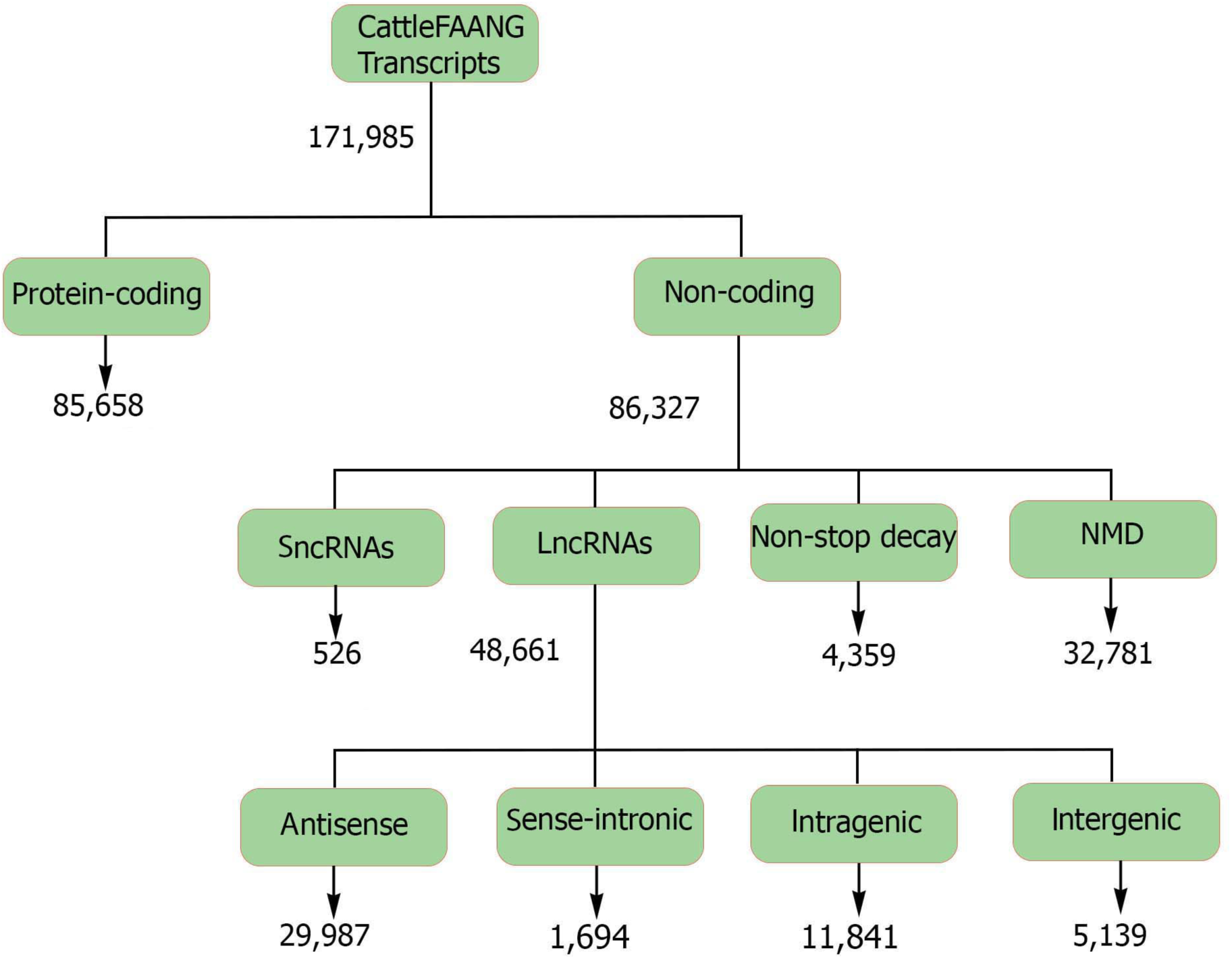
Classification of the predicted transcripts into different biotypes.

There were no significant correlations between the number of RNA-seq reads for a given tissue and the number of unique transcripts identified, except for a modest correlation for the antisense lncRNA class (Supplemental file 1: Fig. S3A). There was a significant positive correlation (p-value 1.3e-04) between the number of unique NMD transcripts in a tissue and the number of protein-coding transcripts, and the NMD transcript class showed the lowest median expression level across tissues, followed by antisense-lncRNAs and sense intronic-lncRNAs (Supplemental file 1: Fig. S2D and Fig. S3B). In addition, there was a significant positive correlation (p-value 3.4e-03) between the number of NMD transcripts and the number of protein-coding transcripts across tissues (Supplemental file 1: Fig. S3A). The expression levels of sncRNAs and protein-coding transcripts were higher (p-values: 1.1e-02 and 2.6e-06, respectively) than that observed for other transcript biotypes (Supplemental file 1: Fig. S2D and Fig. S3B).

### Transcript similarity to other species

Protein/peptide homology analysis of transcripts with coding potential (protein-coding transcripts, lncRNAs, and sncRNAs) revealed a higher conservation rate of protein-coding transcripts (86%) compared to lncRNA and sncRNA transcripts (8%; p-value < 2.2e-16) (Table 2). Bovine non-coding transcripts had significantly (p-value < 2.2e-16) less similarity to other species than protein-coding transcripts (Table 2 and Table 3). Within non-coding transcripts, NSD transcripts showed the lowest conservation rate (35%), followed by sncRNAs (37%), lncRNAs (49%), and NMD transcripts (55%), while sense intronic lncRNAs had the highest conservation rate (60%) compared to other non-coding transcripts (Table 4).

**Table 2.** Protein/peptide homology of transcripts with coding potential

| Transcript biotype | Number of transcripts | Transcripts with protein/peptide homology to other species<sup>1</sup> |
| --- | --- | --- |
| Protein-coding transcripts | 85,658 | 73,268 (86%) |
| sncRNAs and lncRNAs that encode short peptides<sup>2</sup> | 48,425 | 4,054 (8%) |

<sup>1</sup>Number in parentheses indicates the percentage of each transcript biotype.
<sup>2</sup>Open reading frame of 9 to 43 amino acids

**Table 3.**
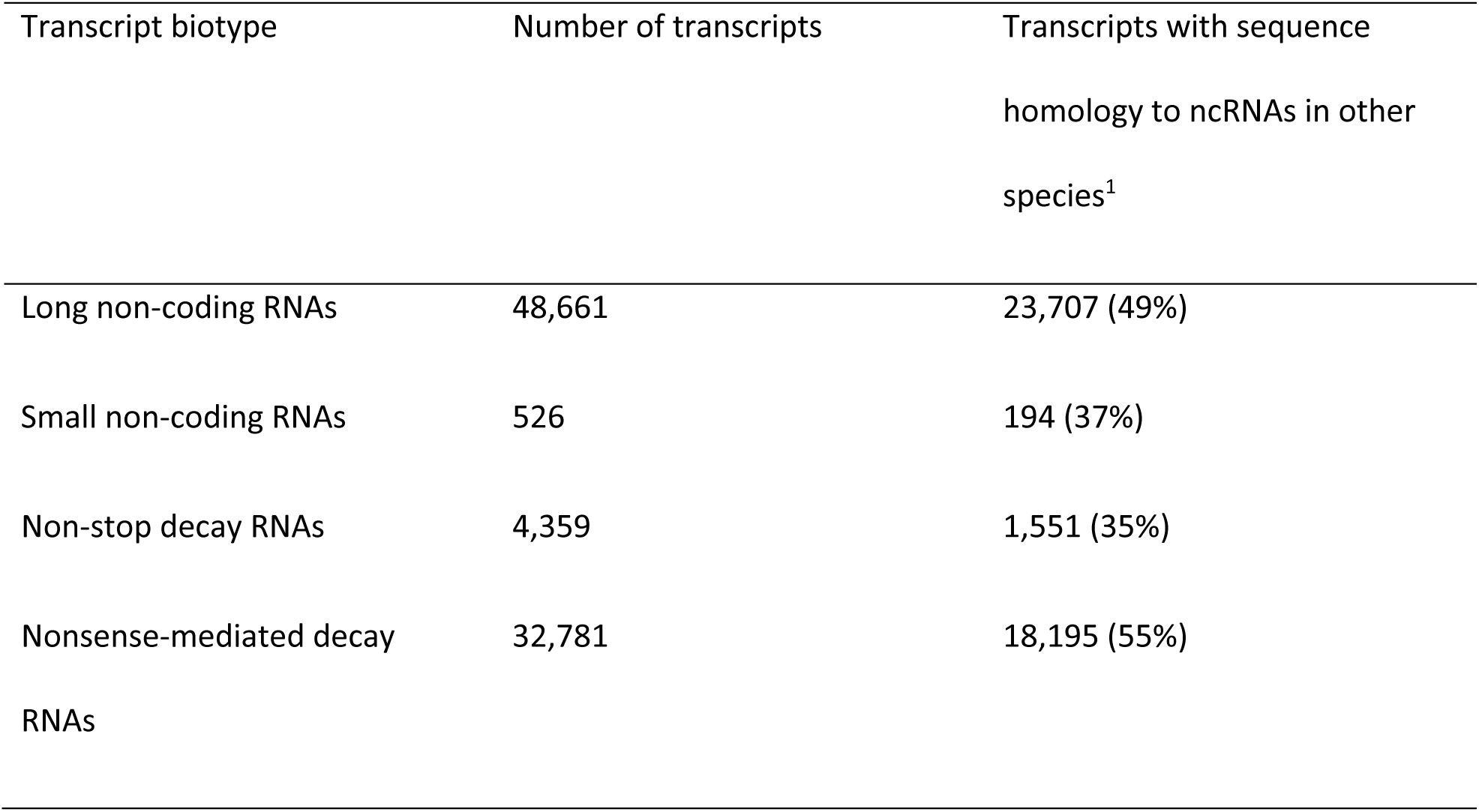
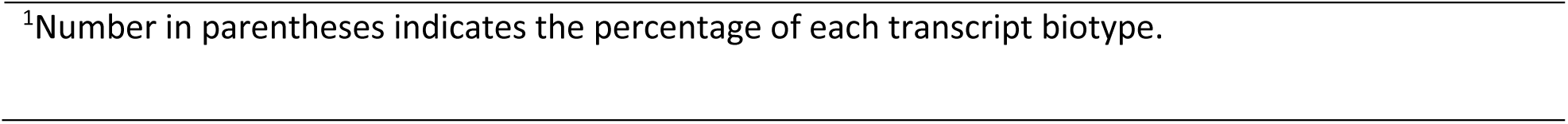
Sequence homology of non-coding transcripts

| Transcript biotype | Number of transcripts | Transcripts with sequence homology to ncRNAs in other species<sup>1</sup> |
| --- | --- | --- |
| Long non-coding RNAs | 48,661 | 23,707 (49%) |
| Small non-coding RNAs | 526 | 194 (37%) |
| Non-stop decay RNAs | 4,359 | 1,551 (35%) |
| Nonsense-mediated decay RNAs | 32,781 | 18,195 (55%) |

<sup>1</sup>Number in parentheses indicates the percentage of each transcript biotype.

**Table 4.** Sequence homology of different types of lncRNAs

| lncRNA biotype | Number of transcripts | Transcripts with sequence homology to ncRNAs in other species<sup>1</sup> |
| --- | --- | --- |
| antisense lncRNAs | 29,987 | 13,793 (46%) |
| sense-intronic lncRNAs | 1,694 | 1,029 (60%) |
| intragenic lncRNAs | 5,569 | 2,314 (41%) |
| intergenic lncRNAs | 11,841 | 5,820 (49%) |

<sup>1</sup>Number in parentheses indicates the percentage of each transcript biotype.

### Transcript diversity across tissues

A median of 70% of protein-coding transcripts were shared between pairs of tissues (Supplemental file 1: Fig. S4A), significantly higher than that was observed for non-coding transcripts (53%; p-value < 2.2e-16; Supplemental file 1: Fig. S5). Clustering of tissues based on protein-coding transcripts was different than that observed based on non-coding transcripts (Supplemental file 1: Fig. S4B and Fig. S5B, Fig. S35F). The fetal tissues clustered together and were generally more similar to one another than to the corresponding adult tissue in both dendrograms, but thymus was closely related to fetal tissues for protein-coding transcript content, while it appeared more similar to lymph nodes, myoblasts, and pregnant/lactating mammary tissue using non-coding transcript profiles. The digestive tract tissues clustered together in the non-coding dendrogram with ileum as a slight outlier, while both jejunum and ileum were distant from the other digestive tissues in the protein-coding transcript profile. The “adult mammary gland” (78 day pregnant) and “virgin mammary gland” samples did not cluster with the three other pregnant/lactating mammary samples nor with each other in either dendrogram. This is mostly likely because: 1) these are from different physiological stages, 2) these were whole tissue samples while the other three pregnant/lactating samples are enriched for mammary gland epithelial cells, 3) the virgin and 78 day pregnant samples are from Hereford background while other pregnant/lactating samples are from Holstein-Frisian breed.

Fetal tissues had significantly higher proportions than adult tissues of unique non-coding transcripts (specifically NSDs, antisense lncRNAs, and intragenic lncRNAs) compared to protein-coding transcripts (p-value < 2.2e-16; Supplemental file 5).

### Transcript validation

Prediction of transcripts and isoforms from RNA-seq data may produce erroneous predicted isoforms. The validity of transcripts was therefore examined by comparison to a library of isoforms taken from Ensembl and NCBI gene sets, plus an assembly produced from all RNA-seq reads, as well as isoforms identified through complete isoform sequencing with Pacific Biosciences and Oxford Nanopore platforms. A total of 118,563 transcripts (70% of predicted transcripts) were structurally validated by at least one other independent dataset. A total of 160,610 transcripts were detected in multiple tissues (96% of predicted transcripts), providing further support for their validity (Fig. 3). All transcripts were also extensively supported by independent data from different technologies such as WTTS-seq, RAMPAGE, histone modification (H3K4me3, H3K4me1, H3K27ac and CTCF), and ATAC-seq (Fig. 3).

**Figure 3.**
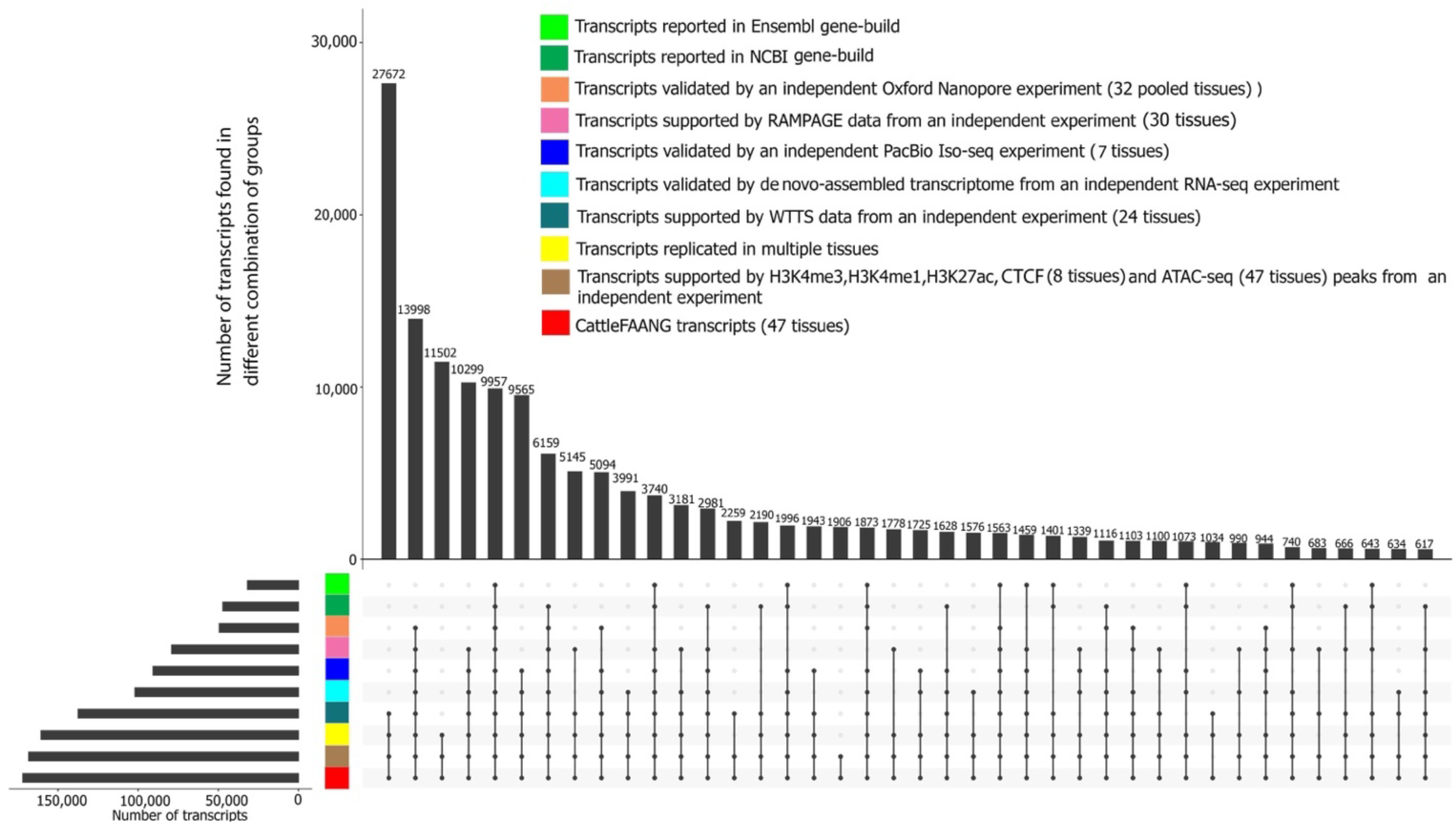
Validation of predicted transcripts using independent data from different technologies.

Comparison of predicted transcript structures with known transcripts in the current bovine genome annotations (Ensembl release 2021-03 and NCBI Release 106) resulted in a total of 52,645 annotated transcripts that exactly matched previously annotated transcripts (31% of all transcripts), including 47,054 annotated NCBI transcripts, 31,740 annotated Ensembl transcripts, and 26,149 transcripts that were common to both annotated gene sets (Fig. 3). The median expression level of known transcripts in their detected tissues (1.8 RPKM) was similar to that observed for novel transcripts (1.4 RPKM) (Supplemental file 1: Fig. S6). Known transcripts were detected in a median of 17 tissues, which was higher (p-value 7.4e-03) than that observed for novel transcripts (median of seven tissues) (Supplemental file 1: Fig. S6). In addition, compared to novel transcripts, annotated transcripts were enriched with protein-coding (p-value 1.37e-02) and spliced transcripts (p-value 3.76e-02).

The median length of coding sequence (CDS) of known transcripts was 1,014 nt, significantly longer than that observed in novel transcripts (510 nt; p-value 0.0) (Additional file1: Fig. S7A). In addition, novel transcripts had longer 5’ UTRs (400 nt) compared to that was observed in known transcripts (300 nt, p-value 2.631E-06; Additional file1: Fig. S7A). Novel transcripts encoding proteins with homology to proteins annotated in other species had longer CDS (687 nt) compared to transcripts without such homology (192 nt; p-value 0.0). Known protein-coding transcripts showed a higher GC content in their 5’ UTRs (61%) than novel transcripts (53%; p-value 5.562E-18), but both classes of transcripts showed similar GC content within their CDS (Supplemental file 1: Fig. S7B).

### Gene level analyses

The transcripts correspond to a total of 35,150 genes, which were detected and classified into protein coding (21,193), non-coding (10,928), and pseudogenes (3,029) (Supplemental file 3 and 4, Fig. 1B, and Fig. 4). The majority of genes detected in each tissue were protein coding (median of 83% of tissue genes), followed by non-coding (median of 14% of tissue genes) and pseudogenes (median of 3% of tissue genes) (Supplemental file 1: Fig. S8). Testis showed the highest number of detected genes with observed transcripts compared to other tissues (Supplemental file 1: Fig. S8). Fetal brain and fetal muscle tissues showed the highest number and percentage of non-coding genes compared to that observed in other tissues (Supplemental file 1: Fig. S8). In addition, more than 40% of transcripts corresponded to non-coding genes (1,271 genes) in fetal brain and fetal muscle. The proportion (6%) and number (1,271) of transcript-producing pseudogenes was higher in testis than in other tissues. There was no significant correlation between the number of input reads and the number of detected genes across tissues, but the numbers of genes from different coding potential classes were significantly correlated across tissues (Supplemental file 1: Fig. S9).

**Figure 4.**
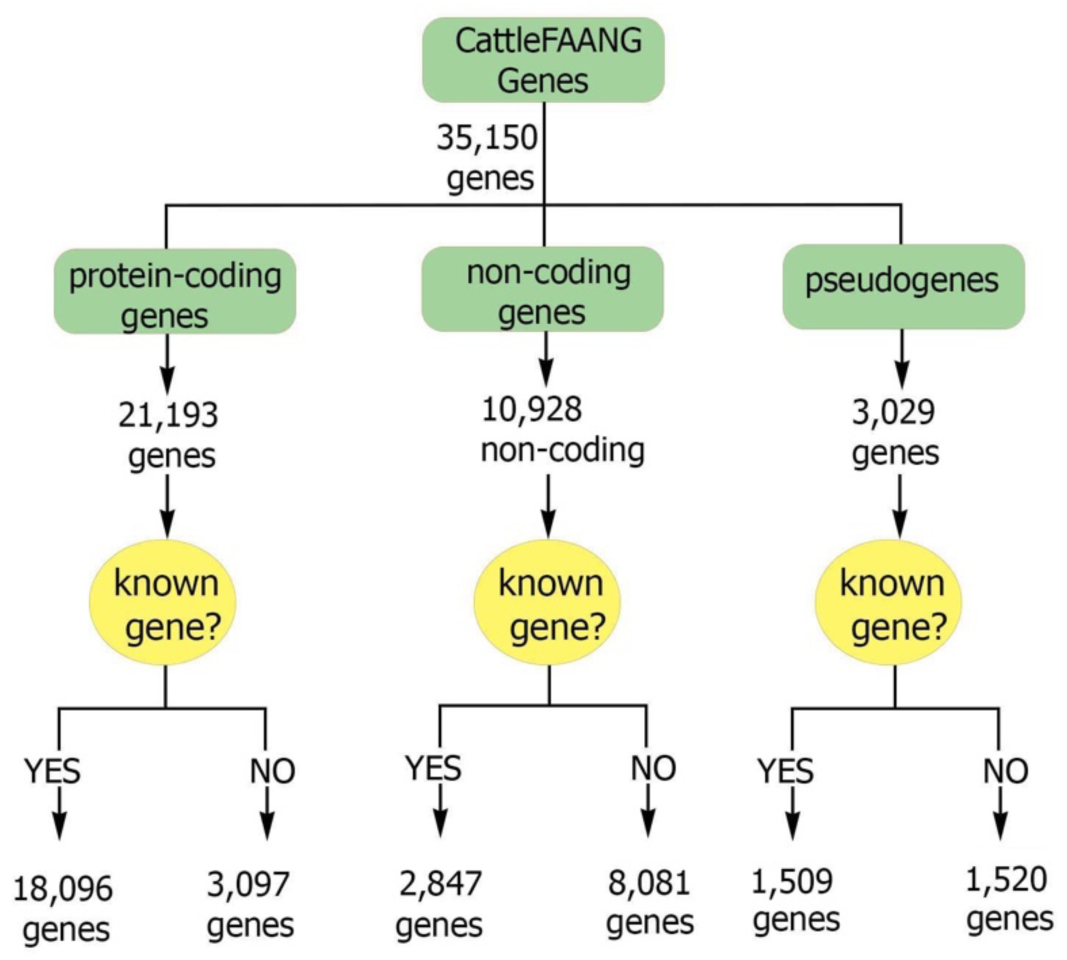
Classification of the predicted genes into different biotypes.

Transcripts corresponding to the predicted genes that had at least one exon overlapping an Ensembl- or NCBI-annotated gene were considered to belong to a known gene. This supported an intersection analysis of predicted and previously annotated genes that indicated 22,452 (64%) of our predicted genes correspond to previously known genes. Approximately 87% of novel transcripts (103,387) were associated with this set of known genes. The remaining 12,698 genes (36% of predicted genes) represent novel genes, i.e., genes not found on Ensembl (release 2021-03) or NCBI (release 106), with which 15% of novel transcripts (22,364 transcripts) were associated. The median number of unique transcripts per known gene (tpg) was four, which was higher than that observed in either the Ensembl (1.5 tpg) or NCBI (2.3 tpg) annotated gene sets, while the median number of transcripts per novel gene was one, with an average of 1.31 and standard deviation of 1.36. Most of the transcripts identified were transcribed from known genes, including 96% of protein-coding transcripts (82,060), 79% of lncRNA transcripts (38,662), 78% of sncRNA transcripts (413), and more than 95% of NMD transcripts (31,422). Known genes were enriched with protein-coding genes (p-value < 2.2e-16). The median transcript abundance from known genes in their detected tissues (6.59 RPKM) was significantly higher than that observed for novel genes (median of 1.68 RPKM; p-value < 2.2e-16; Supplemental file 1: Fig. S10A). The median number of tissues in which known genes were detected (42 tissues) was also significantly higher than that observed for novel genes (median of four tissues; p-value < 2.2e-16; Supplemental file 1: Fig. S10B).

More than a third (37%) of genes with at least one predicted protein-coding transcript displayed either multiple 5’ untranslated regions (UTRs) or multiple 3’ UTRs (median of three 5’ UTRs and three 3’ UTRs per gene) among associated transcript isoforms (Fig. 5). The 496 genes with the highest number of UTRs (the top 5% in this metric) were highly enriched (q-value 1.7E-7) for the “response to protozoan” Biological Process (BP) Gene Ontology (GO) term (Supplemental file 1: Fig. S11 and Supplemental file 6).

**Figure 5.**
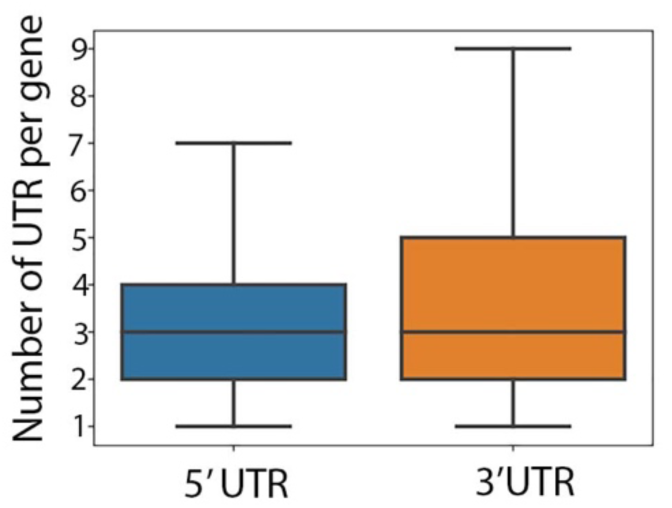
Distribution of the number of 5’ UTRs and 3’ UTRs per gene in genes with multiple UTRs.

A median of 51% of the detected protein-coding genes in each tissue transcribed both protein-coding and non-coding transcripts and were denoted as bifunctional genes. These genes were mostly previously annotated (95%) and had both coding and non-coding transcripts in a median of 21 tissues, representing 57% of their detected tissues (Fig. 6A and B). Protein-coding transcripts and NMD transcripts covered more than 90% of the exonic length in bifunctional genes (Fig. 6C). This percentage was significantly lower for other types of non-coding transcripts transcribed from bifunctional genes (77%, 81%, and 62% for NSD transcripts, sncRNAs, and intragenic lncRNAs, respectively) (Fig. 6C). Although transcript terminal sites (TTS) of transcripts encoded by bifunctional genes were centralized around these genes’ 3’ ends, transcript start sites (TSS) varied greatly among transcript biotypes (Fig. 6C). The TTSs of NSD transcripts, sncRNAs, and intragenic lncRNAs were shifted from their protein-coding genes’ start sites (Fig. 6C). Genes that transcribed both protein-coding and non-coding transcripts in all of their detected tissues (1,661 genes) were highly enriched for “mRNA processing” (q-value 6.08E-16) and “RNA splicing” (q-value 1.35E-14) BP GO terms that were mostly (65%) related to different aspects of transcription and translation (Fig. 6D and Supplemental file 7).

**Figure 6.**
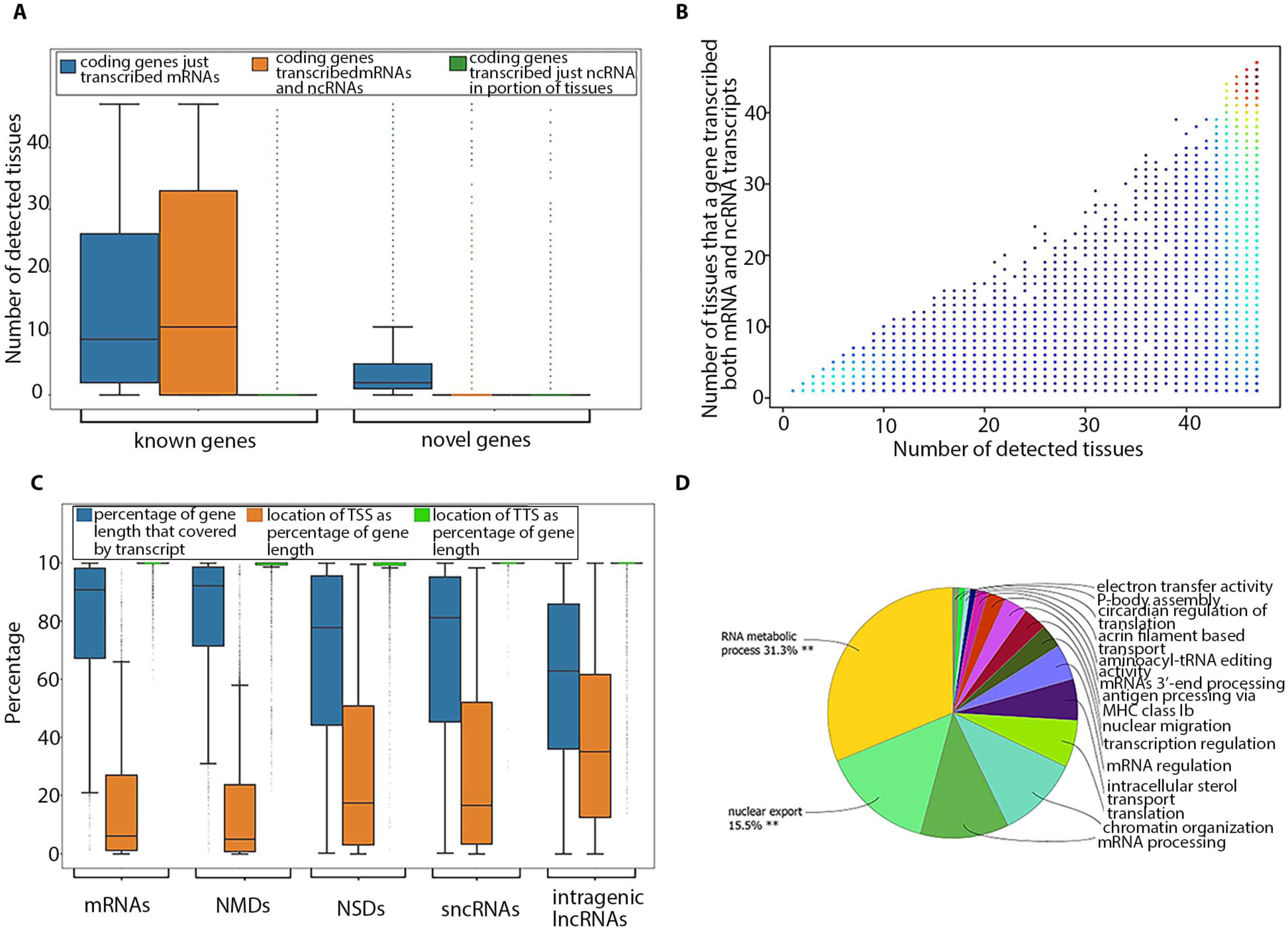
(A) Classification of protein-coding genes based on their novelty and types of encoded transcripts. (B) Number of detected tissues for bifunctional genes. Dots have been color coded based on their density. (C) Location of different transcript biotypes on bifunctional genes. (D) Functional enrichment analysis of genes that remained bifunctional in all of their detected tissues.

A total of 3,744 protein-coding genes (17% of all predicted protein-coding genes) only transcribed non-coding transcripts in a median of two tissues (equivalent to 15% of their detected tissues). Detailed investigation of these genes in tissues from both adult and fetal samples (brain, kidney, muscle, and spleen) revealed the total of 106 non-coding genes (90% known) in fetal tissues that were switched to protein-coding genes with only protein-coding transcripts in their matched adult tissues (Supplemental file 1: Fig. S12). Functional enrichment analysis of these genes resulted in the identification of enriched BP GO terms related to “humoral immune response”, “sphingolipid biosynthetic process”, “negative regulation of wound healing”, “cellular senescence”, “symporter activity”, “regulation of lipid biosynthetic process”, and “filopodium assembly” (Supplemental file 1: Fig. S12, Supplemental file 8).

A median of 32% of protein-coding genes in each tissue expressed at least a single potentially aberrant transcript (PAT), i.e., NMDs and NSDs. In this group of genes, the number of PATs was strongly correlated with the total number of transcripts (median correlation of 0.61 across all tissues). The median expression level of these genes in their detected tissues (11.52 RPKM) was significantly higher (p-value < 2.2e-16) than for protein-coding genes with no PATs (4.48 RPKM). In each tissue, protein-coding genes with PATs showed a significantly higher number of introns (p-value < 2.2e-16; median of 65 introns per gene) than that observed in the remainder of protein-coding genes (median of 15 introns per gene). In addition, genes from this group were detected in a median of 47 tissues, significantly higher (p-value < 2.2e-16) than that observed for the other coding genes (median of 24 tissues), non-coding genes (median of five tissues), and pseudogenes (median of four tissues) (Supplemental file 1: Fig. S13A and B). These genes transcribed a median of two PATs in half (median 54%) of their detected tissues, equivalent to a median of 22% of all their transcripts in each tissue. Protein-coding genes that transcribed PATs as their main transcripts (PATs comprised >50% of their transcripts) in all of their detected tissues were highly enriched with RNA splicing–related BP GO terms (Supplemental file 9).

### Gene similarity to other species

Eighty-five percent of protein-coding genes (18,087) encoded either homologous proteins (17,150 genes or 80% of protein-coding genes) or homologous ncRNAs (7,347 genes or 35% of protein-coding genes) (Supplemental file 1: Fig. S14A). Nineteen percent of protein-coding genes (4,043) encoded cattle-specific proteins (Supplemental file 1: Fig. S14A). The majority of these genes (2,750 or 68%) were either known genes or genes with homology to another cattle gene(s) that has established homology to genes in other species (Supplemental file 1: Fig. S14C). The remaining 32% of cattle-specific, protein-coding genes (1,293 genes or six percent of protein-coding genes) were denoted as protein-coding orphan genes (Supplemental file 1: Fig. S14C). A median of 70 protein-coding orphan genes were detected in each tissue. The expression level of these genes was significantly lower than other types of protein-coding genes (Additional file1: Fig. S15A and B). The median number of detected tissues for protein-coding orphan genes (one tissue) was lower than for other types of protein-coding genes (46 tissues) (Supplemental file 1: Fig. S15C). In addition, protein-coding orphan genes only transcribed protein-coding transcripts in their detected tissue(s).

Fifty percent of non-coding genes (5,559) encoded either homologous short peptides (9-43 amino acids; 5.8% of non-coding genes) or homologous ncRNAs (49% of non-coding genes) (Supplemental file 1: Fig. S14B). There were 5,546 non-coding genes (51% of non-coding genes) that encoded cattle-specific ncRNAs (Supplemental file 1: Fig. S14B). Ninety-nine percent of these genes (5,537 genes) were either known genes or genes with homology to another cattle gene(s) that has established homology to genes in other species (Supplemental file 1: Fig. S14C). The remaining 1% (nine non-coding genes) were denoted as non-coding orphan genes (Supplemental file 1: Fig. S14C). The median number of detected tissues for non-coding orphan genes was 17 tissues, which was higher (p-value < 2.2e-16) than for homologous non-coding genes (six tissues) and protein-coding orphan genes (one tissue) (Supplemental file 1: Fig. S15C).

A total of 3,029 pseudogenes were detected. The median expression level of these genes in their detected tissues was 2.15 RPKM, which was lower than that observed for protein-coding genes (7.08 RPKM) and similar to that observed for non-coding genes (1.7 RPKM) (Supplemental file 1: Fig. S16A). Pseudogenes were detected in a median of four tissues (Supplemental file 1: Fig. S16B). The median number of detected tissues for protein-coding and non-coding genes was 44 tissues and five tissues, respectively (Supplemental file 1: Fig. S16B). In addition, a total of 1,038 pseudogene-derived lncRNAs were detected. The median expression of pseudogene-derived lncRNAs was 1.8 RPKM, similar to that observed for other lncRNAs (1.6 RPKM) (Supplemental file 1: Fig. S17A). In addition, pseudogene-derived lncRNAs were detected in a median of four different tissues, which was lower than observed for other lncRNAs (seven tissues) (Supplemental file 1: Fig. S17B).

Testis had the highest number of detected pseudogene-derived lncRNAs (427), followed by fetal brain (315) (Supplemental file 1: Fig. S8A and B). The correlation between the number of input reads and the number of pseudogene-derived lncRNAs was not significant (0.25, p-value 0.09).

### Gene diversity across tissues

Tissue similarities increased dramatically from transcript level to gene level (Supplemental file 1: Fig. S4A, Fig. S5A, Fig. S18A, Fig. S19A). The median percentage of shared genes between pairs of tissues was significantly higher in protein-coding genes compared to non-coding genes (90% and 57%, respectively; p-value < 2.2e-16; Supplemental file 1: Fig. S18A, Fig. S19A). Clustering of tissues based on protein-coding genes was similar to that observed based on protein-coding transcripts (Supplemental file 1: Fig. S18B, Fig. S19B). The same result was observed in non-coding genes and transcripts. In addition, clustering of tissues based on protein-coding genes was different than that of non-coding genes (Supplemental file 1: Fig. S4B, Fig. S5B, Fig. S18B, Fig. S19B, Fig. S35F).

Tissues with both fetal and adult samples (brain, kidney, muscle, and spleen) were used to investigate gene biotype differences between these developmental stages. Similar to what was observed at transcript level, fetal tissues were significantly enriched for non-coding genes and pseudogenes and were depleted for protein-coding genes (p-value < 2.2e-16; Supplemental file 10). These results were consistent across all tissues with both adult and fetal samples (Supplemental file 10).

### Gene validation

A total of 32,460 genes (92% of predicted genes) were structurally validated by independent datasets (PacBio Iso-Seq data, ONT-seq data, *de novo* assembled transcripts from RNA-seq data, or Ensembl and NCBI gene sets). In addition, a total of 31,635 genes (90% of predicted genes) were detected in multiple tissues (31,635 genes or 90%) (Fig. 7). All genes were extensively supported by independent data from different technologies such as WTTS-seq, RAMPAGE, histone modification (H3K4me3, H3K4me1, H3K27ac) and CTCF-DNA binding, and ATAC-seq data generated from the samples.

**Figure 7.**
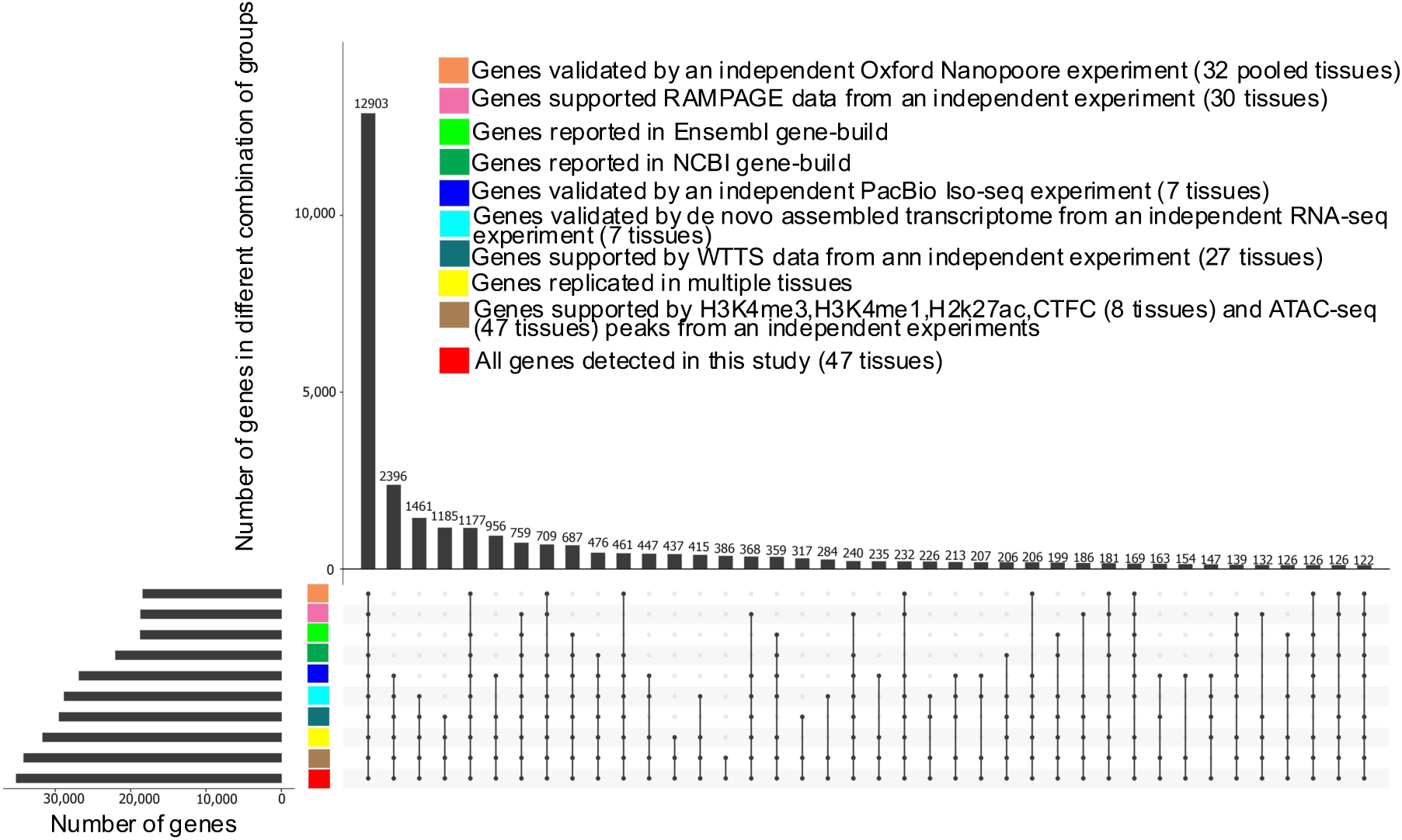
Validation of predicted genes using independent data from different technologies

### Identification and validation of known gene border extensions

This new bovine gene set annotation extended (5ʹ end extension, 3ʹ end extension, or both) more than 11,000 known Ensembl or NCBI gene borders. Extensions were longer on the 3ʹ side, but the median increase was 104 nucleotides (nt) for the 5’ end (Table 5). To validate gene border extensions, independent WTTS-seq (24 tissues) and RAMPAGE datasets (30 tissues) were utilized. More than 80% of known gene border extensions were validated by independent data (Fig. 8). The extension of known gene borders on both ends resulted in an approximate nine-fold expression increase of these genes in the new bovine gene set annotation compared to their matched Ensembl and NCBI genes (Table 6). This effect was smaller in known genes extended only on 5ʹ or 3ʹ ends (Table 6).

**Table 5.** Gene border extensions in current ARS-UCD1.2 genome annotations by *de novo* assembled transcriptome from short-read RNA-seq data

| Annotation | Type of gene extension | Number of genes | Median extension<br>(nucleotides) |
| --- | --- | --- | --- |
| Ensembl<br>(release 2021-03) | 5' extension only | 1,848 | 128 |
|  | 3' extension only | 5,701 | 422 |
|  | Both ends extended | 4,874 | 122, 5'<br>439, 3' |
| NCBI<br>(Release 106) | 5' extension only | 2,214 | 80 |
|  | 3' extension only | 5,496 | 126 |
|  | Both ends extended | 3,613 | 66, 5'<br>210, 3' |

**Figure 8.**
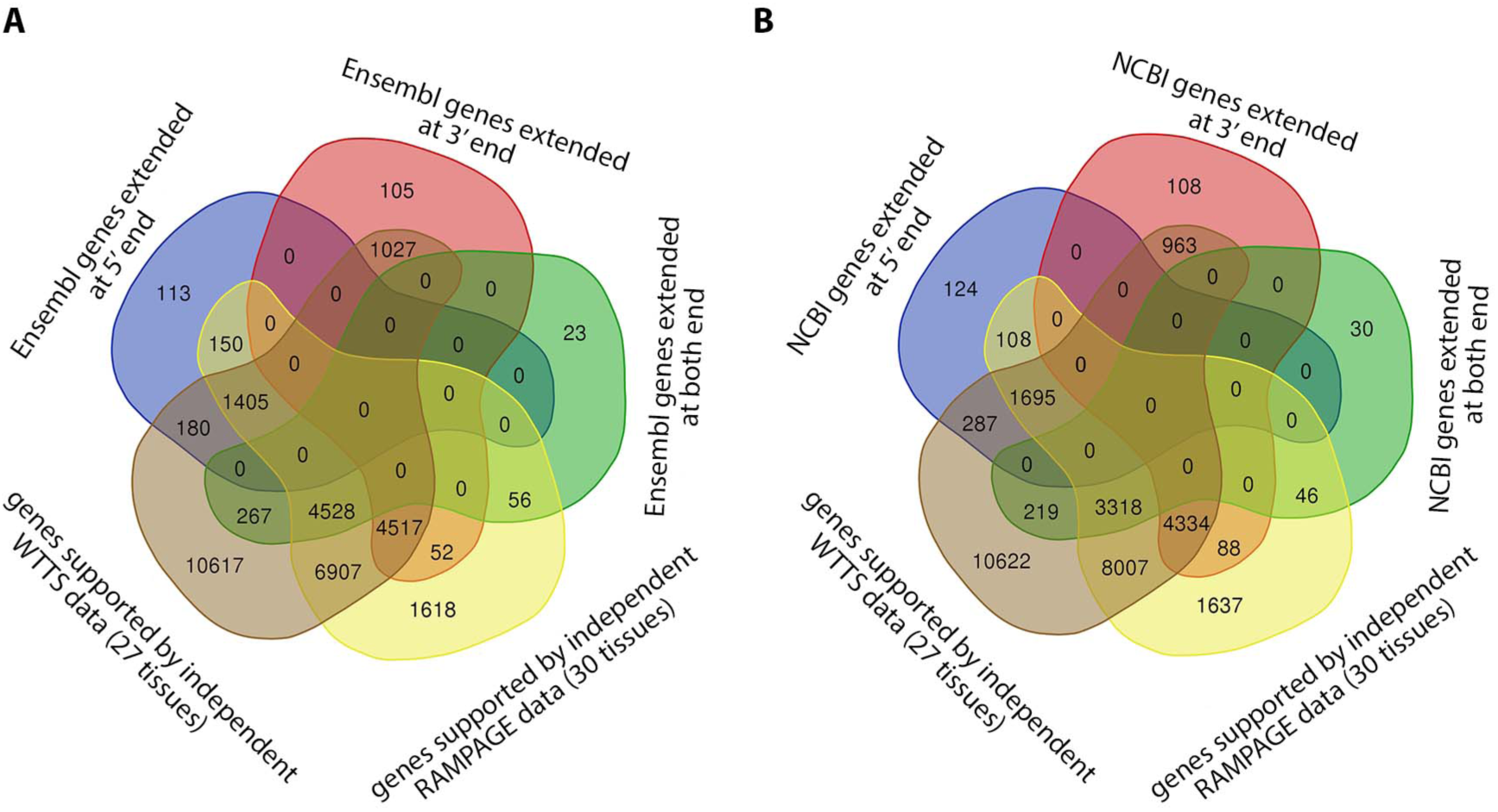
Functional enrichment analysis of non-coding genes in fetal tissues that were switched to protein coding with only coding transcripts in their matched adult tissue

**Table 6.**
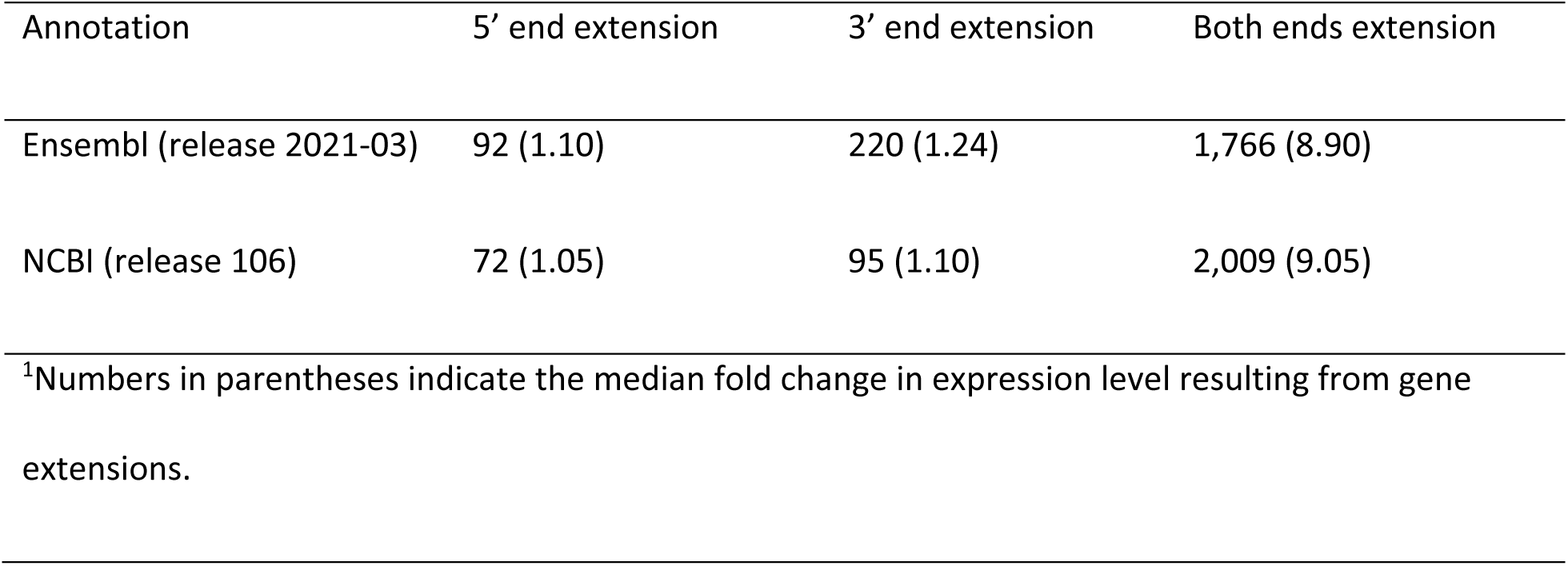
Median number of reads mapped to the extended region of known genes^1^

### Alternative splicing events

Alternative splicing (AS) events (Supplemental file 1: Fig. S20A) are commonly distinguished in terms of whether RNA transcripts differ by inclusion or exclusion of an exon, in which case the exon involved is referred to as a “skipped exon” (SE) or “cassette exon”, “alternative first exon”, or “alternative last exon”. Alternatively, spliced transcripts may also differ in the usage of a 5’ splice site or 3’ splice site, giving rise to alternative 5’ splice site exons (A5Es) or alternative 3’ splice site exons (A3Es), respectively. A sixth type of alternative splicing is referred to as “mutually exclusive exons” (MXEs), in which one of two exons is retained in RNA but not both. However, these types are not necessarily mutually exclusive; for example, an exon can have both an alternative 5’ splice site and an alternative 3’ splice site, or have an alternative 5’ splice site or 3’ splice site but be skipped in other transcripts. A seventh type of alternative splicing is “intron retention”, in which two transcripts differ by the presence of an unspliced intron in one transcript that is absent in the other. An eighth type of alternative splicing is “unique splice site exons” (USEs), in which two exons overlap with no shared splice junction. A total of 102,502 bovine transcripts (80% of spliced transcripts) were involved in different types of AS events, a large increase over NCBI (73,423 transcripts) and Ensembl (37,299 transcripts) annotations (Additional file1: FigureS20B). Skipped exons were observed in a greater number of transcripts compared to other types of AS events (Supplemental file 1: Fig. S21).

A median of 60% of tissue transcripts showed at least one type of AS event (Supplemental file 1: Fig. S22A). There was no significant correlation between the number of input reads and the number of AS event transcripts across tissues (Supplemental file 1: Fig. S22B).

The median expression level of AS transcripts (111,366 transcripts or 65% of transcripts) was 1.38 RPKM, similar to that observed for other types of transcripts (1.58RPKM) (Supplemental file 1: Fig. S23A). In addition, AS transcripts were detected in a median of 10 tissues (Supplemental file 1: Fig. S23B), which was higher than for the other transcript types (median of nine tissues). AS transcripts were enriched with protein-coding transcripts (p-value < 2.2e-16). Meanwhile, transcripts that did not show AS events, i.e., unspliced transcripts and spliced transcripts from single transcript genes, were enriched for non-coding transcripts (p-value < 2.2e-16). A median of 67% of protein-coding genes showed at least one type of AS event. In contrast, this was only 3% in non-coding genes. In most cases, AS events did not change transcript biotypes (Supplemental file 1: Fig. S24). In addition, a switch from protein-coding to ncRNAs was the main biotype change resulting from AS events (Supplemental file 1: Fig. S24).

A median of four AS events were detected in alternatively spliced genes (14,260 genes or 40% of genes) (Supplemental file 1: Fig. S25). The top five percent of genes with the highest number of AS events (2,734 genes, Fig. 35A) were highly enriched for several BP GO terms related to different aspects of RNA splicing (Supplemental file 1: Fig. S26B, Supplemental file 11).

Comparison of tissues with both fetal and adult samples (brain, kidney, Latissimus Dorsi (LD) muscle, and spleen) revealed a significantly higher rate of AS events in fetal tissues (only genes expressed in both fetal and adult samples were included in this analysis) (Supplemental file 1: Fig. S27).

### Tissue specificity

Nine percent of all genes (3,174) and transcripts (15,562) were only detected in a single tissue and were denoted as tissue-specific (Supplemental file 1: Fig. S28A). The majority of tissue-specific genes (75%) and transcripts (84%) were novel. Forty-nine percent of tissue-specific transcripts (11,748) were produced by known genes. The majority of tissue-specific genes (61%) and transcripts (57%) were protein-coding (Supplemental file 1: Fig. S28A and B). In addition, more than 70% of tissue-specific transcripts (11,222) were transcribed from non-tissue-specific genes. Compared to other tissues, testis and thymus had the highest number of tissue-specific genes and transcripts (Supplemental file 1: Fig. S28C, Supplemental file 12). The expression level of tissue-specific genes and transcripts was significantly lower than that of their non-tissue-specific counterparts (p-value < 2.2e-16; Supplemental file 1: Fig. S28D). A median of 71% of tissue-specific transcripts showed any type of AS event in their detected tissues (Supplemental file 1: Fig. S29). This was only 3.9% for tissue-specific genes (Supplemental file 1: Fig. S29). Testis, myoblasts, mammary gland, and thymus had the highest proportion of tissue-specific genes displaying any type of AS event (Supplemental file 1: Fig. S29).

A total of 16,806 multi-tissue detected genes (53% of all multi-tissue detected genes) and 74,487 multi-tissue detected transcripts (51% of all multi-tissue detected transcripts) showed Tissue Specificity Index (TSI) scores (Supplemental file 13) greater than 0.9 and were expressed in a tissue-specific manner. These genes and transcripts were detected in a median of six tissues and four tissues, respectively (Supplemental file 1: Fig. S30A and B). Functional enrichment analysis of the top five percent of genes with the highest TSI score (3,171 genes) resulted in the identification of “sexual reproduction” (p-value 3.06e-24) and “fertilization” (p-value 1.04e-8) as their top enriched BP GO terms (Supplemental file 1: Fig. S30C-E, Supplemental file 14).

### Tying genes to phenotypes

There were 9,800 predicted genes identified as the closest gene to an existing QTL (QTL-associated genes) in their detected tissues (Supplemental file 15). These genes had either QTLs located inside (6,511 genes) or outside (5,306 genes) their genomic borders (either from their 5’ end or 3’ end) with a median distance of 51.9 kilobases (KB) and a maximum distance of 2.6 million bases (MB) (Supplemental file 1: Fig. S31). The majority of QTL-associated genes were known genes (8,130 genes or 83%). In addition, the median number of AS events in these genes (eight) was significantly higher than that observed in other genes (median of seven AS events; p-value 5.69e-09).

### Potential testis-pituitary axis

Testis tissue was not clustered with any other tissues and had the highest number of tissue-specific genes (1,195 genes) compared to the rest of the tissues (Supplemental file 1: Fig. S4, Fig. S5, Fig. S18, and Fig. S19). Testis-specific genes were highly enriched with different traits related to fertility (e.g., percentage of normal sperm and scrotal circumference), body weight (e.g., body weight gain and carcass weight), and feed efficiency (e.g., residual feed intake) (Supplemental file 16). The extent of testis-pituitary axis involvement in the “percentage of normal sperm” was investigated using animals with both testis and pituitary samples (three samples per tissue). The *SPACA5* gene was the only testis-specific gene with a (or gene encoding a) signal peptide (SP) that was close to the “percentage of normal sperm” QTLs. The expression of this gene in testis samples showed significant positive correlation with 70 pituitary genes that were closest to the “percentage of normal sperm” QTLs (Supplemental file 1: Fig. S32, Supplemental file 17). These pituitary genes were enriched with the “signal transduction in response to DNA damage” BP GO term (Supplemental file 1: Fig. S32). In addition, the expression of testis genes that encoded signal peptide that were close to the “percentage of normal sperm” QTLs was significantly correlated with expression of pituitary genes close to this trait (Fig. 9, Supplemental file 18). The same result was observed for the pituitary-testis tissue axis (Supplemental file 1: Fig. S33, Supplemental file 19).

**Figure 9.**
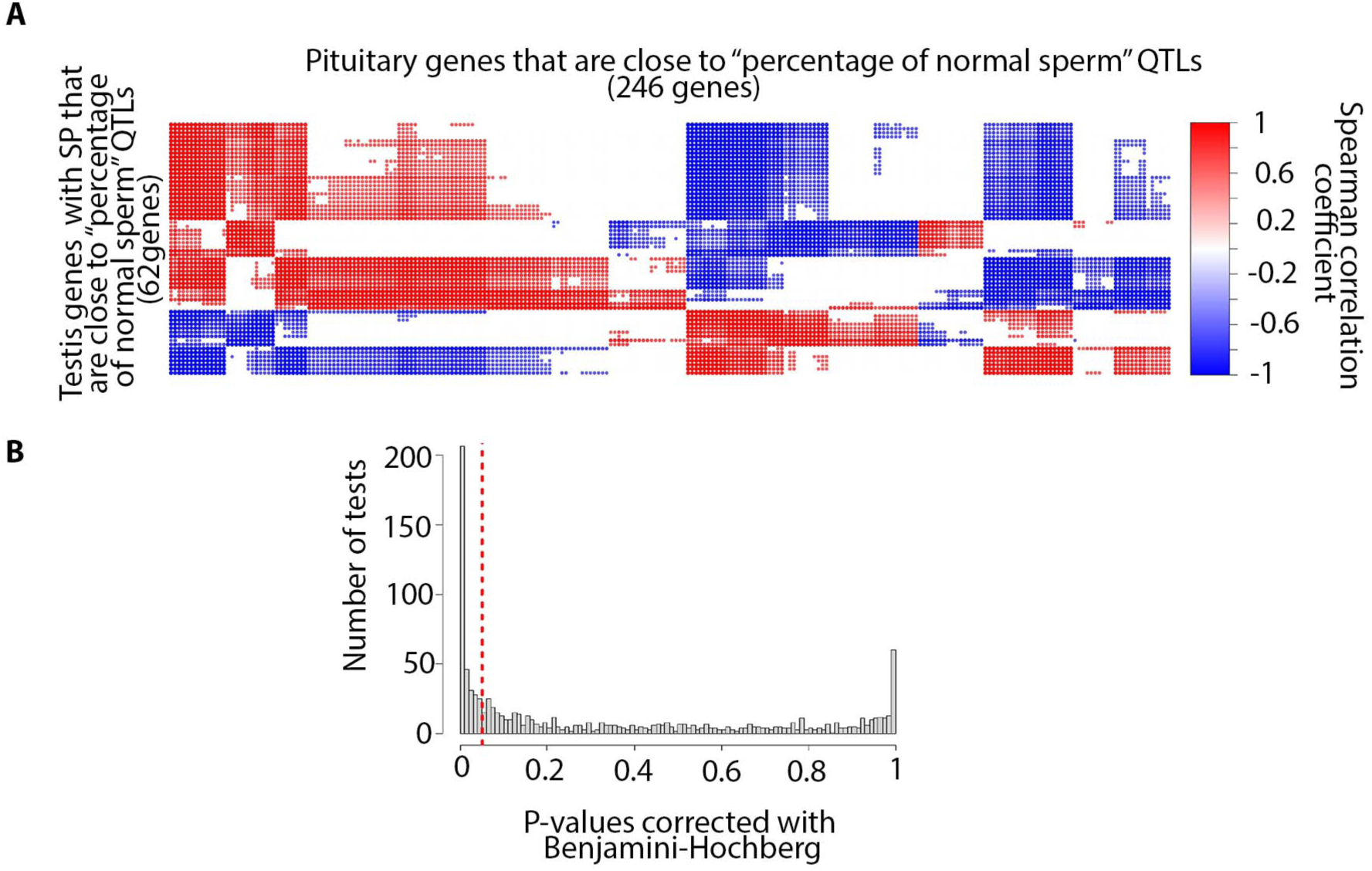
(A) Correlation between testis genes with signal peptides that were close to the “percentage of normal sperm” QTL and pituitary genes closest to this trait (reference correlations). (B) Distribution of p-values resulting from a right-sided t-test between reference correlation coefficients and correlation coefficients derived from random chance (see methods for details).

### Trait similarity network

The extent of genetic similarity between different bovine traits was investigated using their associated QTLs. A total of 1,857 significantly similar trait pairs (184 different traits) were identified and used to create a bovine trait similarity network (https://www.animalgenome.org/host/reecylab/a; Supplemental file 20).

### miRNAs

A total of 2,007 miRNAs (at least ten mapped reads in each tissue) comprised of 973 known and 1,034 novel miRNAs were detected (Supplemental file 21). In each tissue, a median of 704 known miRNAs and 549 novel miRNAs were detected (Fig. 10A). The median expression of novel miRNAs was 0.10 Reads Per Million (RPM), which was significantly lower than that observed for known miRNAs (0.41 RPM; p-value 3.25e-25; Fig. 10B). In addition, novel miRNAs were detected in a median of 23 tissues, significantly lower than for known miRNAs (43 tissues; p-value 1.00e-45; Fig. 10C). A median of 84.53% of miRNAs were shared between pairs of tissues (Supplemental file 1: Fig. S34). Clustering of tissues based on miRNAs was similar to what was observed based on non-coding genes (Supplemental file 1: Fig. S35).

**Figure 10.**
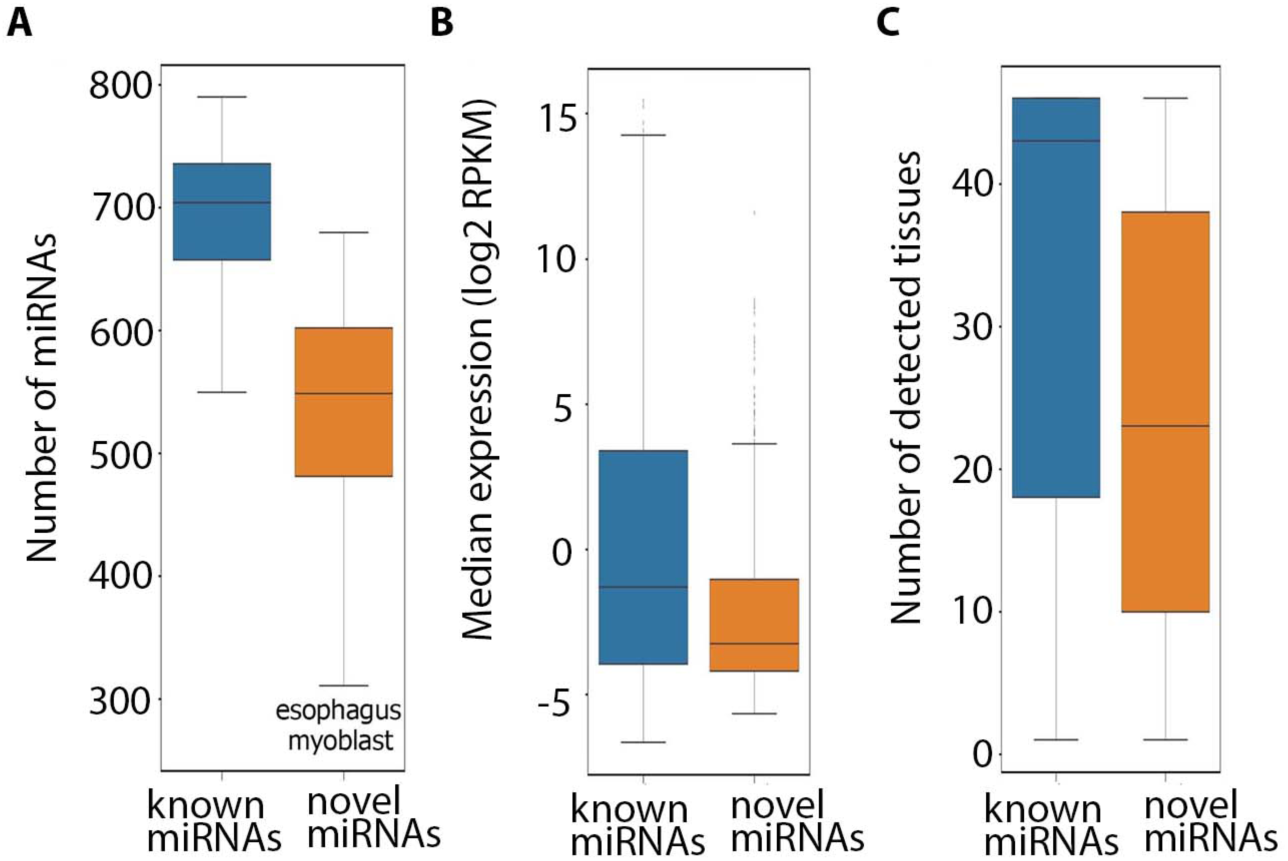
(A) Distribution of the number of detected known and novel miRNAs across tissues. (B) Expression of known and novel miRNAs across their detected tissues. (C) Number of detected tissues for known and novel miRNAs.

A total of 113 miRNAs (5.6%) were detected in a single tissue and were denoted as tissue-specific (Supplemental file 1: Fig. S36A). The proportion of tissue-specific miRNAs was higher for novel miRNAs, such that 75% of the tissue-specific miRNAs (85) were novel. The number of novel miRNAs was higher in pre-adipocytes compared to other tissues, followed by fetal gonad and testis (Supplemental file 1: Fig. S36B). Novel miRNAs showed a significantly lower expression level compared to known miRNAs (p-value 1.4e-19; Supplemental file 1: FigureS36 C). In addition, a total of 1,047 multi-tissue detected miRNAs (55% of all multi-tissue detected miRNAs) had a TSI score greater than 0.9 and were expressed in a tissue-specific manner (Additional file1: Fig. S36D). These miRNAs were detected in a median of 19 tissues (Supplemental file 1: Fig. S36E).

Chromatin features across 500-base pair (bp) windows surrounding upstream of miRNA precursors’ start sites or downstream of miRNA precursors’ terminal sites from independent cattle experiments were used to investigate the relationship between miRNAs and chromatin accessibility. More than 99% of novel miRNAs (1,027) and 94% of known miRNAs (923) were supported by at least one of the H3K4me3, H3K4me1, H3K27ac, CTCF-DNA binding, or ATAC-seq peaks (Fig. 11).

### Summary of detected transcripts, genes, and miRNAs

The numbers of detected transcripts, genes, and miRNAs in different tissues are summarized in Supplemental file 1: Fig. S37. In addition, the number of known and novel genes, transcripts, and miRNAs in different tissues are summarized in Supplemental file 1: Fig. S38.

**Figure 11.**
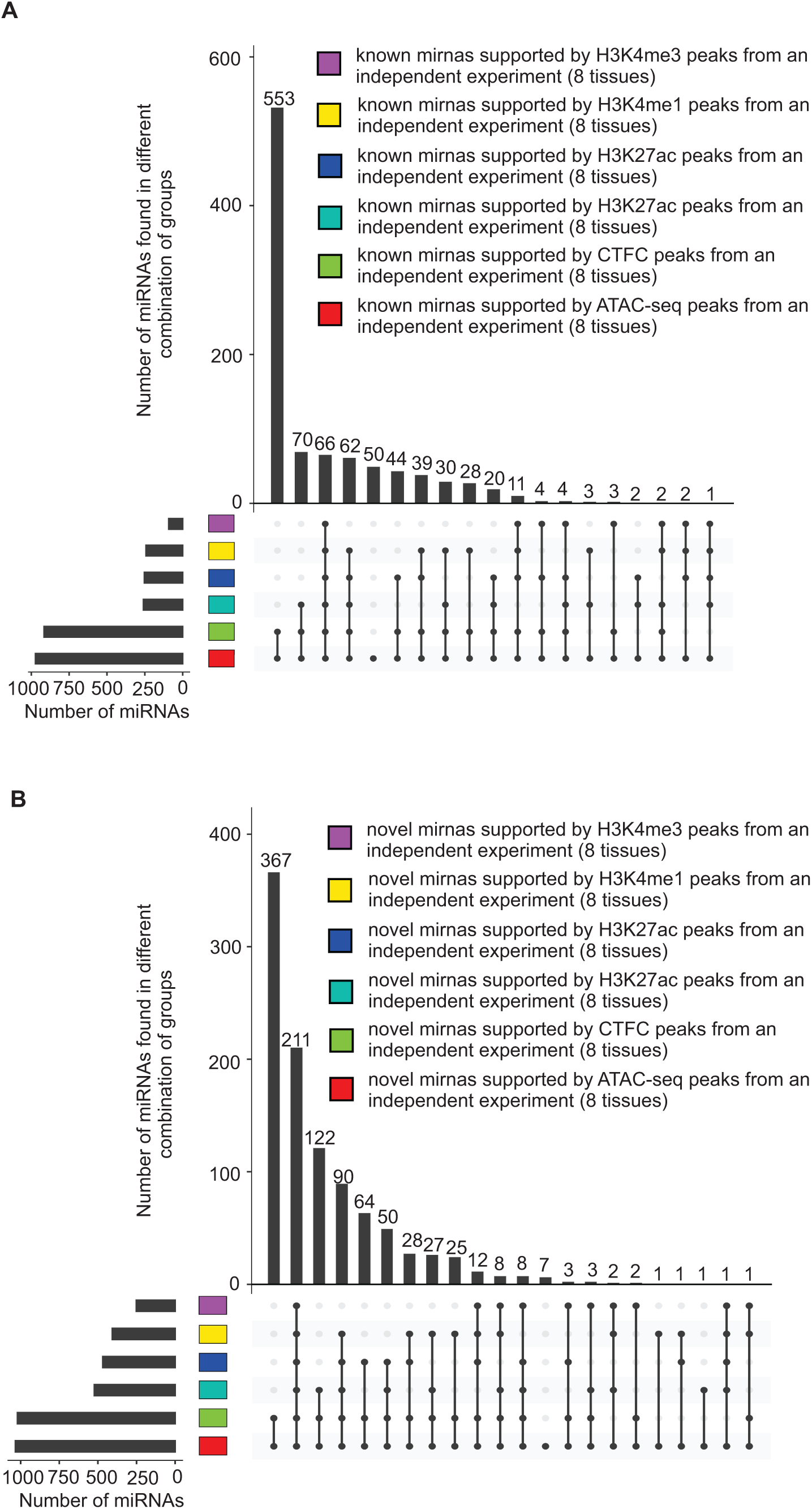
Validation of detected known (A) and novel (B) miRNAs using different histone mark data

## Discussion

Despite many improvements in the current bovine genome annotation ARS-UCD1.2 assembly (Ensembl release 2021-03 and NCBI release 106) compared to the previous genome assembly (UMD3.1), these annotations are still far from complete (Goszczynski et al. 2021; Halstead et al. 2021). In this study, using RNA-seq and miRNA-seq data from 47 different bovine tissues/cell types, 12,698 novel genes and 1,034 novel miRNAs were identified that have not been reported in current bovine genome annotations (Ensembl release 2021-03, NCBI release 106 and miRbase (Kozomara et al. 2019)). In addition, we identified protein-coding transcripts with a median ORF length of 270 nt for 822 known bovine genes that have been annotated as non-coding in current bovine genome annotations (Supplemental file 1: Fig. S14C). The high frequency of validation of these novel genes and novel miRNAs using multiple independent datasets from different technologies verifies the improvement in terms of the number of genes and miRNAs using our methods.

Five prime and 3‘untranslated region length plays a critical role in regulation of mRNA stability, translation, and localization (Jereb et al. 2018). However, only a single 5’ UTR and 3’ UTR per gene is annotated in current bovine genome annotations (Ensembl release 2021-03 and NCBI release 106), and variations in UTR length are not available. In this study, 7,909 genes (22% of predicted genes) with multiple UTRs were identified. Genes with multiple 5ʹ UTRs are common, primarily due to the presence of multiple promoters (Araujo et al. 2012) or alternative splicing mechanisms within 5’ UTRs (Araujo et al. 2012). Fifty-four percent of human genes have multiple transcription start sites (Araujo et al. 2012). In addition, the length of 3ʹ UTRs often varies within a given gene, due to the use of different poly-adenylation (polyA) sites (Jereb et al. 2018; Gerber et al. 2021).

In this study, around 50% of detected protein-coding genes in each tissue transcribed both coding and non-coding transcript isoforms. Several studies have shown evidence of the existence of bifunctional genes with coding and non-coding potential using high-throughput RNA sequencing (RNA-seq) and ribosome footprinting followed by sequencing (Ribo-seq) (Andrews and Rothnagel 2014; Kumari and Sampath 2015; Nam et al. 2016). More than 20% of human protein-coding genes have been reported to transcribe non-coding isoforms, often generated by alternative splicing (Gonzàlez-Porta et al. 2013) and recurrently expressed across tissues and cell lines (Nam et al. 2016). A considerable number of non-coding isoform variants of protein-coding genes appear to be sufficiently stable to have functional roles in cells (Mayba et al. 2014). It has been shown that the proportion of non-coding isoforms from protein-coding genes dramatically increases during myogenic differentiation of primary human satellite cells and decreases in myotonic dystrophy muscles (Hubé et al. 2011). In this study, 106 non-coding genes were identified in fetal tissues that switched to protein-coding genes in their matched adult tissues. Taken together this supports the notion that protein-coding/non-coding transcript switching plays an important role in tissue development in cattle as well.

Nonsense-mediated RNA decay is an evolutionarily conserved process involved in RNA quality control and gene regulatory mechanisms (Kurosaki et al. 2019). For instance, the RNA-binding protein polypyrimidine tract binding protein 1 (*PTBP1*) can promote the transcription of NMD transcripts via alternative splicing, which negatively regulates its own expression (Wollerton et al. 2004). In this study, NMD transcripts comprised 19% of bovine transcripts that were transcribed from 30% of bovine genes (10,498). In humans, NMD-mediated degradation can affect up to 25% of transcripts (Nickless et al. 2017) and 53% of genes (Supek et al. 2021). As expected, in this study, the majority of genes that transcribed NMD transcripts were protein coding (83% or 8,687 genes), while a considerable portion (17%) were pseudogenes. Many pseudogenes are known to give rise to NMD transcripts (Mitrovich and Anderson 2005; Colombo et al. 2017). Bioinformatic study of the human transcriptome revealed that 78% of NMD transcript–producing genes were protein coding, followed by pseudogenes (nine percent), long intergenic noncoding RNAs (six percent), and antisense transcripts (four percent) (Colombo et al. 2017).

Despite the important regulatory function of lncRNAs and miRNAs, very low numbers of these elements have been annotated in the current bovine genome annotations (Table 7). In this study, a total of 10,789 lncRNA genes and 2,007 miRNA genes were detected in the bovine transcriptome, which is similar to what has been reported for the human transcriptome (Table 7). While, a total of 3,770 human miRNAs and 1,203 cattle miRNAs have been reported in miRbase (Kozomara et al. 2019). Small non-coding RNAs detected from RNA-seq data did not overlap with known or novel miRNA precursors or known sncRNAs reported in Ensembl or NCBI gene builds.

**Table 7.** Comparison of different gene builds based on gene biotypes

| Species | Gene build | Protein-coding genes | lncRNA genes | miRNA genes | Other types of small non-coding genes<sup>1</sup> | Pseudo-genes |
| --- | --- | --- | --- | --- | --- | --- |
| Bovine<br>(ARS-UCD1.2) | Ensembl | 21,880 | 1,480 | 951 | 2,209 | 492 |
|  | (Release 2021-03) |  |  |  |  |  |
|  | NCBI | 21,039 | 5,179 | 797 | 3,249 | 4,569 |
|  | (Release 106) |  |  |  |  |  |
|  | Cattle FAANG<sup>2</sup> | 21,193 | 10,789 | 2,007 | 139 | 3,029 |
|  |  | (18,096) | (2,847) | (973) | (0) | (1,509) |
| Human<br>(GRCh38.104) | Ensembl | 20,442 | 16,876 | 1,877 | 2,930 | 15,266 |
|  | (release 2021-03) |  |  |  |  |  |

<sup>1</sup>Small nucleolar RNAs, small nuclear RNAs, small Cajal body specific RNAs, small conditional RNAs, and tRNAs
<sup>2</sup>Numbers in parentheses indicate the number of novel RNAs in each biotype.

In this study, 1,038 pseudogene-derived lncRNAs were identified that were recurrently expressed across tissues and cell types. Ever-increasing evidence from different studies suggests pseudogene-expressed RNAs are key components of lncRNAs (Milligan and Lipovich 2014; Stewart et al. 2019; Lou et al. 2020). lncRNAs expressed from pseudogenes have been shown to regulate genes with which they have sequence homology (Milligan and Lipovich 2014; Stewart et al. 2019) or to coordinate development and disease in metazoan systems (Milligan and Lipovich 2014).

Correct annotation of gene borders has an important role in defining promoter and regulatory regions. Our novel transcriptome analysis extended (5ʹ-end extension, 3ʹ-end extension, or both) more than 11,000 known Ensembl or NCBI gene borders. Extensions were longer on the 3ʹ side, which was relatively similar to that we observed in the pig transcriptome using PacBio Iso-Seq data (Beiki et al. 2019).

A growing body of evidence indicates that a considerably large portion of lncRNAs encode microproteins that are less conserved than canonical Open Reading Frames (ORFs) (Anderson et al. 2015; Mackowiak et al. 2015; Olexiouk et al. 2016; Li and Liu 2019; Wei and Guo 2020). In this study, a vast majority (98%) of predicted lncRNAs had short ORFs (<44 amino acids) that were less conserved than canonical ORFs (Table 2).

Alternative splicing is the key mechanism to increase the diversity of the mRNA expressed from the genome and is therefore essential for response to diverse environments. In this study, skipped exons and retained introns were the most prevalent AS events identified in the bovine transcriptome, similar to what has been observed in other vertebrates and invertebrates (Sammeth et al. 2008). A higher rate of AS events was observed in fetal tissues compared to their adult tissue counterparts. The same result has been observed in a recently published study in humans (Mazin et al. 2021).

We hypothesized that the integration of the gene/transcript data with previously published QTL/gene association data would allow for the identification of potential molecular mechanisms responsible for a) tissue-tissue communication as well as b) genetic correlations between traits. To test the first hypothesis, we developed a novel approach to study the involvement of tissue-tissue interconnection in different traits based on the integration of the transcriptome with publicly available QTL data. In particular, the interconnection between testis and pituitary tissues with respect to the “percentage of normal sperm” trait was investigated in more detail. This resulted in the identification of the regulation of ubiquitin-dependent protein catabolic process, the regulation of Nuclear factor-κB (NF-κB) transcription factor activity, and Rab protein signal transduction as key components of this tissue-tissue interaction (Supplemental file 18 and 19). Interestingly, genes that were closest to “percentage of normal sperm” QTLs, and also encoded signal peptides in both testis and pituitary tissues, were highly enriched for the BP GO term “regulation of ubiquitin-dependent protein catabolic process” (Supplemental file 18 and 19). The expression of these genes in testis tissue was significantly correlated with expression levels of pituitary genes closest to “percentage of normal sperm” QTLs that were highly enriched for the “positive regulation of NF-kappaB transcription factor activity” BP GO term (Supplemental file 1: Fig. S32 and Supplemental file 18). Activation of NF-κB requires ubiquitination, and this modification is highly conserved across different species (Chen and Chen 2013). NF-κB induces secretion of adrenocorticotropic hormone from the pituitary (Karalis et al. 2004), which directly stimulates testosterone production by the testis (O’Shaughnessy et al. 2003). In addition, ubiquitinated proteins in testis cells are required for the progression of mature spermatozoa (Richburg et al. 2014). The expression levels of pituitary genes closest to “percentage of normal sperm” QTLs that also encoded signal peptides were significantly correlated with expression levels of testis genes closest to “percentage of normal sperm” QTLs (Supplemental file 1: Fig. S33). These testis genes were highly enriched for the “Rab protein signal transduction” BP GO term (Supplemental file 19). Rab proteins have been reported to be involved in male germ cell development (Kumar et al. 2016). Thus, it appears that integration of gene data with QTL/association data can be used to identify putative molecular pathways underlying tissue-tissue communication mechanisms.

To test the second hypothesis, we also developed a novel approach to study trait similarities based on the integration of the transcriptome with publicly available QTL data. Using this approach, we could identify significant similarity between 184 different bovine traits. For example, clinical mastitis showed significant similarity with 23 different cattle traits that were greatly supported by published studies, such as milk yield (Rajala-Schultz et al. 1999), milk composition traits (Martí De Olives et al. 2013), somatic cell score (Halasa and Kirkeby 2020), foot traits (Remnant et al. 2019), udder traits (Miles et al. 2019), daughter pregnancy rate (Lima et al. 2020), length of productive life (Hertl et al. 2018) and net merit (Kaniyamattam et al. 2020). Similar results were observed for residual feed intake, which showed significant similarity with 14 different traits such as average daily feed intake (Green et al. 2013), average daily gain (Elolimy et al. 2018), carcass weight (Weber et al. 2013), feed conversion ratio (Yi et al. 2018), metabolic body weight (Liu and VandeHaar 2020), subcutaneous fat (Clare et al. 2018), and dry matter intake (Houlahan et al. 2021).

Taken together, these results identify a list of candidate genes that might harbor genetic variation responsible for the genetic mechanisms underlying genetic correlations (Supplemental file 18 and 1. If this is the case, in the future, these novel methods should be able to predict the impact of a given set of genetic variants that are associated with a trait of interest on other traits that were not measured in a given study. This might then lead to the optimization of variants used (or not used) in genomic selection to minimize any non-beneficial effect of selection on selected traits.

## Conclusions

In-depth analysis of multi-omics data from 47 different bovine tissues/cell types provided evidence to improve the annotation of thousands of protein-coding, lncRNA, and miRNA genes. These validated results increase the complexity of the bovine transcriptome (number of transcripts per gene, number of UTRs per gene, lncRNA transcripts, AS events, and miRNAs), comparable to that reported for the highly annotated human genome. We provided direct evidence that the predicted novel transcripts extend existing known gene models, by verifying such extensions using independent WTTS-seq and RAMPAGE data. We utilized a novel approach to integrate the transcriptome with publicly available QTL data and showed its application in a study of tissue axis involvement in different traits and genetic similarity between different traits. This approach is particularly important in the selection of indicator traits for breeding purposes, study of artificial selection side effects in livestock species, and functional annotation of poorly annotated livestock genomes.

## Methods

### Tissue and cell collection, total RNA extraction and construction of RNA-seq, miRNA-seq, WTTS-seq, ATAC-seq, and ChIP-seq libraries

#### Cell sample collections

Skeletal muscle and subcutaneous fat samples were collected from Angus-crossbred steers slaughtered at the Virginia Tech Meat Center. Satellite cells were isolated from skeletal muscle by pronase digestion as described previously (Leng et al. 2019). The isolated satellite cells were activated to proliferate as myoblasts by culturing in growth medium composed of Dulbecco’s Modified Eagle Medium (DMEM), 10% fetal bovine serum (FBS), and 1% antibiotics-antimycotics. To induce myoblasts to differentiate into myocytes, myoblasts cultured in growth medium were switched to differentiation medium composed of DMEM and 2% horse serum for 2 days. Preadipocytes from subcutaneous fat were isolated by collagenase digestion as previously described (Hausman et al. 2008). To induce preadipocytes to differentiate into adipocytes, preadipocytes were initially cultured in growth medium (DMEM/F12, 10% FBS, 1% antibiotics-antimycotics) to reach confluency, then in induction medium (DMEM/F12, 10% FBS, 1% antibiotics-antimycotics, 10 μg/mL insulin, 1 μM dexamethasone, 0.5 mM isobutyl methylxanthine, and 200 μM indomethacin) for 2 days, and lastly in maintenance medium (DMEM/F12, 10% FBS, 1% antibiotics-antimycotics, 1 μg/mL insulin) for 10 days.

#### Adult tissue collections

Procedures for tissue collection followed the Animal Care and Use protocol (#18464) approved by the Institutional Animal Care and Use Committee (IACUC), University of California, Davis (UCD). Four cattle (2 males and 2 females) were slaughtered at UCD using captive bolt under USDA inspection at 14 months old and were intact male and female Line 1 Herefords that had the same sire, provided by Fort Keogh Livestock and Range Research Lab (Tixier-Boichard et al. 2021). Tissue samples were flash frozen in liquid nitrogen then stored at –80 °C until further assay processing.

#### Fetal tissue collections

Fetal sample collection and tissue collection were approved by the Institutional Animal Care and Use Committee (IACUC), University of Idaho (2017-67). Four pregnant females at day 78 of gestation Line 1 Herefords were slaughtered at UI meats lab using captive bolt under USDA inspection. Animals were provided by Fort Keogh Livestock and Range Research Lab (Tixier-Boichard et al. 2021). Tissue samples were flash frozen in liquid nitrogen then stored at –80 °C until further assay processing.

#### RNA-seq library construction

Tissue samples (Supplemental file 22) were collected from storage at −80 °C and ground to a powder using a mortar and pestle and liquid nitrogen. The tissue was next homogenized in QIAzol Lysis Reagent (Qiagen Catalog No. 79306) using a QIAshredder spin column (Qiagen Catalog No. 79656). After centrifugation, the lysate was mixed with chloroform, shaken vigorously for 15 sec, incubated for 2 – 3 min at room temperature, and centrifuged for 15 min at 12,000 x g at 4°C. The upper, aqueous phase was transferred to a new collection tube and 1.5 vol of 100% ethanol was added and mixed thoroughly by pipetting up and down several times. Total RNA was then isolated from the sample using the RNeasy Mini Kit (Qiagen Catalog No. 74106) according to the manufacturer’s instructions. Contaminating DNA was removed by treating total RNA with DNase (AM1906, Ambion). Total RNA quantity was measured with the Quant-It RiboGreen RNA Assay Kit (Life Technologies Corp., Carlsbad, CA) and quality assessed by fragment analysis (Advance Analytical Technologies, Inc., Ankeny IA).

#### Mammary gland tissue collection and RNA-seq library construction

The 16 animals used in this study were Holstein-Friesian heifers from a single herd managed at the AgResearch Research Station in Ruakura, NZ. All experimental protocols were approved by the AgResearch, NZ, ethics committee, and carried out according to their guidelines. Samples were collected from the same animals at 5 time points: virgin state before pregnancy between 13 and 15 months of age (virgin), mid-pregnant at day 100 of pregnancy, late pregnant ∼2 weeks pre-calving, early lactation ∼2 weeks post-calving, and at peak lactation, 34-38 days post-calving. Tissue samples were obtained by mammary biopsy using the Farr method (Farr et al. 1996). Lactating cows were milked before biopsy and sampled within 5 hours of milking. Biopsy sites were clipped and given aseptic skin preparation (povidone iodine base scrub and iodine tincture) and subcutaneous local anesthetic (4 ml per biopsy site). Core biopsies were taken using a powered sampling cannula (4.5 mm internal diameter) inserted into a 2 cm incision. The resulting samples of mammary gland parenchyma measured 70 mm in length, with a 4 mm diameter. Small slices from each sample were preserved for histology before mammary epithelial organoids were separated from surrounding adipose and connective tissue to allow for secretory-specific signals in the RNA-seq analysis. In preparation for isolating organoids, tissue samples were digested in a freshly prepared collagenase solution containing 0.2% collagenase A (Roche), 0.05% trypsin (1:250 powder, 100U/ml Gibco), hyaluronidase (Sigma), 5% fetal calf serum (Hyclone), Pen/Strep/Fungizone solution (Hyclone) or 5 µg/ml Gentamycin (Sigma) in DMEM/F12 (Gibco) with 10 ng/ml insulin. Samples were minced to a fine slurry and incubated in this freshly prepared collagenase solution (10 ml solution/g tissue) for 3.5 hours at 37°C in a 50 ml conical tube with slow shaking (120 rpm). Digested tissue was centrifuged at 453 x g for 10 min at 4°C, after which the supernatant and fat layers were discarded, and the pellet was gently resuspended in 5 ml DMEM/F12 without serum. A further 5 ml DMEM/F12 without serum was added, and the sample was centrifuged at 453 x g for 10 min at 4°C. The media was discarded, and the pellet was gently resuspended in 10 ml DMEM/F12 and centrifuged for another 10 min at 453 x g and 4°C. The media was discarded, and pellet resuspended in 10 ml DMEM/F12 for a third time, and the sample centrifuged in a series of brief spins achieved by allowing the centrifuge to reach 453 x g for two seconds before applying the brake. These brief pulse spins were repeated at least 4 times, or until examining the sample under a microscope revealed primarily epithelial organoid clusters and very few single cells. At this point, the organoid pellet was resuspended in 1 ml TRIzol and stored at −80°C until RNA isolation. High-quality total RNA (RIN > 7) was extracted from frozen mammary epithelial organoid pellets using NucleoSpin® miRNA isolation kit (MACHEREY-NAGEL) according to the manufacturer’s protocol, isolating large and small (<200 nt) fractions separately. The “large” RNA fraction was used to prepare strand-specific poly-A+ RNA-seq libraries for sequencing. The “small” RNA fraction was used to make miRNA-seq libraries using NEXTflex^TM^ Small RNA-Seq Kit v3.

#### miRNA-seq library construction

Tissue samples (Supplemental file 22) were collected similarly to the method described in the previous section. QIAseq miRNA Library Kit (Qiagen, cat no. 331505) and QIAseq miRNA NGS 96 Index IL Kit (Qiagen, cat no. 331565) were used to isolate miRNAs from all tissues except mammary gland. miRNAs from mammary gland were isolated using NEXTflex^TM^ Small RNA-Seq Kit v3 (Illumina) according to the manufacturer’s instructions. The isolated miRNA was subjected to 3’ ligation to ligate a pre-adenylated DNA adaptor to the 3’ ends of all miRNAs. An RNA adaptor was then ligated to the 5’ end of the mature miRNA to complete 5’ ligation. cDNA synthesis was completed using a reverse transcriptase (RT) primer containing integrated unique molecular identifiers (UMI). The RT primer bound to the 3’ adaptor region and facilitated conversion of the 3’/5’ ligated miRNAs into cDNA while a UMI was assigned to every miRNA molecule. After reverse transcription, a clean-up of the cDNA was performed using a streamlined magnetic bead-based method. Library amplification was accomplished by a universal forward primer from a plate being paired with 1 of 96 dried reverse primers in the same plate (Qiagen, cat no. 331565) to assign each sample a unique custom index. Following library amplification, a clean-up of the miRNA library was performed using a streamlined magnetic bead-based method. Libraries were then evaluated for quantity and quality measures before being normalized and pooled for llumina sequencing (1×50bp).

#### WTTS-seq library construction

Construction of the WTTS-seq libraries from tissue samples (Supplemental file 22) involved fragmentation, polyA+ RNA enrichment, first-strand cDNA synthesis by reverse transcription and second-strand cDNA synthesis by PCR as described previously (Zhou et al. 2016). The starting material was 2.5 µg of total RNA per library, which was fragmented with 1 μl of 10X RNA fragmentation buffer (Ambion, AM8740), followed by enrichment of polyA+ RNA using Dynabeads (Ambion 61002). The polyA+ RNA molecules were then used for the first-strand cDNA synthesis with both 5’ adaptor (switching primer, 100 µM) and 3’ adaptor (containing oligo (dT10), 100 µM) catalyzed by the SuperScript III reverse transcriptase (200 U/μl) (Invitrogen, 18080). The first-strand cDNA molecules were chemically enriched with RNases I and H and used to synthesize the second-strand cDNA using PCR. Base PCR conditions were as follow: initial denaturation at 98 °C for 30 s, PCR cycles of 98 °C for 10 s, 50°C for 30 s, and 72°C for 30 s, and final extension at 72°C for 10 min. The size-selected cDNA (200 – 500 bp) was purified with SPRI beads (Agencourt AMPure XP beads, Beckman Coulter, Brea, CA) and sequenced using an Ion PGM™ Sequencer at Washington State University.

#### ChIP-seq library construction

ChIP-seq experiments (H3K4me3, H3K4me1, H3K27ac and H3K27me3) were performed on flash-frozen tissue samples (Supplemental file 22) using the iDeal ChIP-seq kit (Diagenode Cat. #C01010059, Denville, NJ). Briefly, 20–30 mg powdered tissue was cross-linked with 1% formaldehyde for 8 min and quenched with 100 μl of glycine for 10 min. Nuclei were obtained by centrifugation at 2000 × g for 5 min and resuspended in 600 μl of iS1 buffer for incubation on ice for 30 min. Chromatin was sheared using the Bioruptor Pico between 10 and 15 cycles depending on the tissues. For immunoprecipitation experiments, ∼1–1.5 μg of sheared chromatin was used as input with 1 μg (histone modifications) or 1.5 μg (CTCF) of antibody following the protocol from the kit. The following antibodies used were from Diagenode: H3K4me3 (comes with Diagenode iDeal Histone kit), H3K27me3 (#C15410069), H3K27ac (#C15410174), H3K4me1 (#C15410037), and CTCF (#15410210). An input (no antibody) was performed for each sample. Libraries were constructed using the NEBNext Ultra DNA library prep kit (New England Biolabs #E7645L, Ipswich, MA). Libraries were sequenced on the Illumina HiSeq 4000 platform, generating 50 bp single-end reads.

#### ATAC-seq library construction

Frozen tissue samples (Supplemental file 22) were pulverized under liquid nitrogen using mortar and pestle. Permeabilized nuclei were obtained by resuspending pulverized tissue (5-15 mg) in 250 µL Nuclear Permeabilization Buffer (0.2% IGEPAL-CA630 [I8896, Sigma], 1 mM DTT [D9779, Sigma], Protease inhibitor [05056489001, Roche], and 5% BSA [A7906, Sigma] in PBS [10010-23, Thermo Fisher Scientific]), and incubating for 10 min on a rotator at 4°C. Nuclei were then pelleted by centrifugation for 5 min at 500 x g at 4°C. The pellet was resuspended in 25 µL ice-cold Tagmentation Buffer (33 mM Tris-acetate [pH = 7.8; BP-152, Thermo Fisher Scientific], 66 mM K-acetate [P5708, Sigma], 11 mM Mg-acetate [M2545, Sigma], 16% DMF [DX1730, EMD Millipore] in molecular biology grade water [46000-CM, Corning]). An aliquot was then taken and counted by hemocytometer to determine nuclei concentration. Approximately 50,000 nuclei were resuspended in 20 µL ice-cold Tagmentation Buffer and incubated with 1 µL Tagmentation enzyme (FC-121-1030, Illumina) at 37 °C for 30 min with shaking at 500 rpm. The tagmentated DNA was purified using MinElute PCR purification kit (28004, Qiagen). The libraries were amplified using NEBNext High-Fidelity 2X PCR Master Mix (M0541, NEB) with primer extension at 72°C for 5 min, denaturation at 98°C for 30 s, followed by 8 cycles of denaturation at 98°C for 10 s, annealing at 63°C for 30 s and extension at 72°C for 60 s. Amplified libraries were then purified using MinElute PCR purification kit (28004, Qiagen), and two size selection steps were performed using SPRIselect bead (B23317, Beckman Coulter) at 0.55X and 1.5X bead-to-sample volume ratios, respectively. ATAC-seq libraries were sequenced on an Illumina Nextseq 500 platform using Nextra V2 sequencing chemistry to generate 2 × 75 paired-end reads.

### Sequencing the transcriptomes of seven bovine tissues by using the PacBio Iso-Seq and Illumina RNA-Seq technologies

Frozen tissue samples (Supplemental file 22) were pulverized by grinding with disposable mortar and pestle in liquid nitrogen. RNA was extracted using TRIzol reagent as directed by the manufacturer (Invitrogen) with integrity examined using a BioAnalyzer (Agilent). Only samples with RIN values >8 were used for cDNA synthesis. Libraries for RNA-seq short-read sequencing were prepared using the TruSeq RNA Kit following the “TruSeq RNA Sample Preparation v2 Guide” as recommended by the manufacturer (Illumina). RNA-seq libraries were sequenced on a NextSeq500 instrument. IsoSeq libraries for long-read sequencing were prepared using the SMRTbell Template Prep Kit 1.0. First strand cDNA synthesis was performed with approximately 1 µg of extracted RNA from each tissue using the Clontech SMARTer PCR cDNA Synthesis Kit (Clontech) as directed by the manufacturer. cDNA was then converted to SMRTbell template library following the “Iso-Seq using Clontech cDNA Synthesis and BluePippin Size Selection” protocol as directed by the manufacturer (Pacific Biosciences). Three size fraction pools for each tissue were prepared using the BluePippin instrument (Sage Science), representing insert sizes of 1-2 kb, 2-3 kb, and 3-6 kb. The two smaller fractions were sequenced in three to five SMRT cells on an RSII instrument (Pacific Biosciences), and the largest fraction sequenced in five or six cells, using P6/C4 chemistry. The sequences were processed into HQ isoforms using SMRT Analysis v6.0 for each tissue independently but with all size fractions within tissue included in the analysis.

### RNA-seq data analysis and transcriptome assembly

Single-end Illumina RNA-Seq reads (75 bp) from each tissue sample were trimmed to remove the adaptor sequences and low-quality bases using Trim Galore (version 0.6.4) (Krueger 2019) with --quality 20 and --length 20 option settings. The resulting reads were aligned against ARS-UCD1.2 bovine genome using STAR (version 020201) (Dobin et al. 2013) with a cut-off of 95% identity and 90% coverage. FeatureCounts (version 2.0.2) (Liao et al. 2014) was used to quantify genes reported in the NCBI gene build (version 1.21) with -Q 255 -s 2 --ignoreDup --minOverlap 5 option settings. The resulting gene counts were adjusted for library size and converted to Counts Per Million (CPM) values using SVA R package (version 3.30.0) (Leek et al. 2021). In each tissue, sample similarities were checked using hierarchical clustering and regression analysis of gene expression values (log2 based CPM), and outlier samples were detected and removed from downstream analysis. Samples from each tissue were combined to get the most comprehensive set of data in each tissue. To reduce the processing time due to huge sequencing depth, the trimmed reads were in silico normalized using insilico_read_normalization.pl from Trinity package (version 2.6.6) (Grabherr et al. 2011) with -- JM 350G and --max_cov 50 option settings. Normalized RNA-seq reads were aligned against ARS-UCD1.2 bovine genome using STAR (version 020201) (Dobin et al. 2013) with a cut-off of 95% identity and 90% coverage. The normalized reads were assembled using *de novo* Trinity software (version 2.6.6) (Grabherr et al. 2011) combined with massively parallelized computing using HPCgridRunner (v1.0.1) (Hass 2015) and GNU parallel software (Tange 2018). The resulting transcript reads were collapsed and grouped into putative gene models (clustering transcripts that had at least a one-nucleotide overlap) by the pbtranscript-ToFU from SMRT Analysis software (v2.3.0) (PacificBiosciences 2018) with min-identity = 95%, min-coverage = 90% and max_fuzzy_junction = 15 bp, whereas the 5ʹ-end and 3’-end difference were not considered when collapsing the reads. Base coverage of the resulting transcripts was calculated using mosdepth (version 0.2.5) (Pedersen and Quinlan 2018). Predicted transcripts were required to have a minimum of three times base coverage in their detected tissues. The predicted acceptor and donor splice sites were required to be canonical and supported by Illumina-seq reads that spanned the splice junction with 5-nt overhang. Spliced transcripts with the exact same splice junctions as their reference transcripts but that contained retained introns were removed from analysis, as they were likely pre-RNA sequences. Unspliced transcripts with a stretch of at least 20 A’s (allowing one mismatch) in a genomic window covering 30 bp downstream of their putative terminal site were removed from analysis, as they were likely gDNA contaminations. To decrease the false positive rate, unspliced transcripts that were only detected in a single tissue were removed from downstream analysis. The resulting transcripts from each tissue were re-grouped into gene models using an in-house Python script. The collapsed transcripts from the different tissues were then merged using an in-house Python script to create the RNA-seq based bovine transcriptome.

The resulting transcripts and genes were quantified using align_and_estimate_abundance.pl from the Trinity package (version 2.6.6) (Grabherr et al. 2011) with --aln_method bowtie -- est_method RSEM --SS_lib_type RF option settings.

“Isoform” and “transcript” terms are used interchangeably throughout the manuscript.

### PacBio Iso-Seq data analysis

PacBio Iso-Seq data has been processed as described for the pig transcriptome (Beiki et al. 2019) with the following exceptions. Errors in the full-length, non-chimeric (FLNC) cDNA reads were corrected with the preprocessed RNA-Seq reads from the same tissue samples using the combination of proovread (v2.12) (Hackl et al. 2014) and FMLRC (v1.0.0) (Wang et al. 2018) software packages. Error rates were computed as the sum of the numbers of bases of insertions, deletions, and substitutions in the aligned FLCN error-corrected reads divided by the length of aligned regions for each read (Table 8).

The RNA-seq-based transcriptome was assembled as described in the previous section.

**Table 8.**
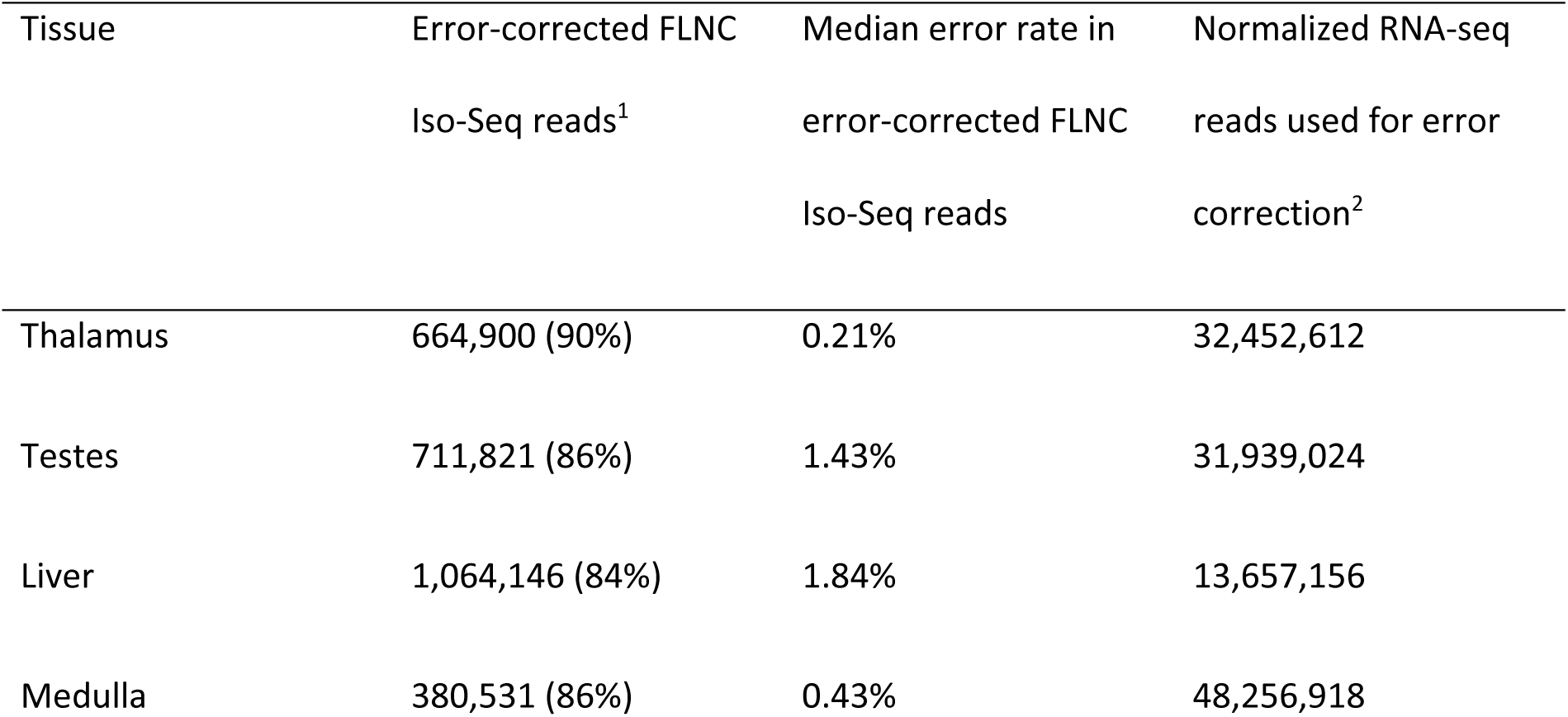
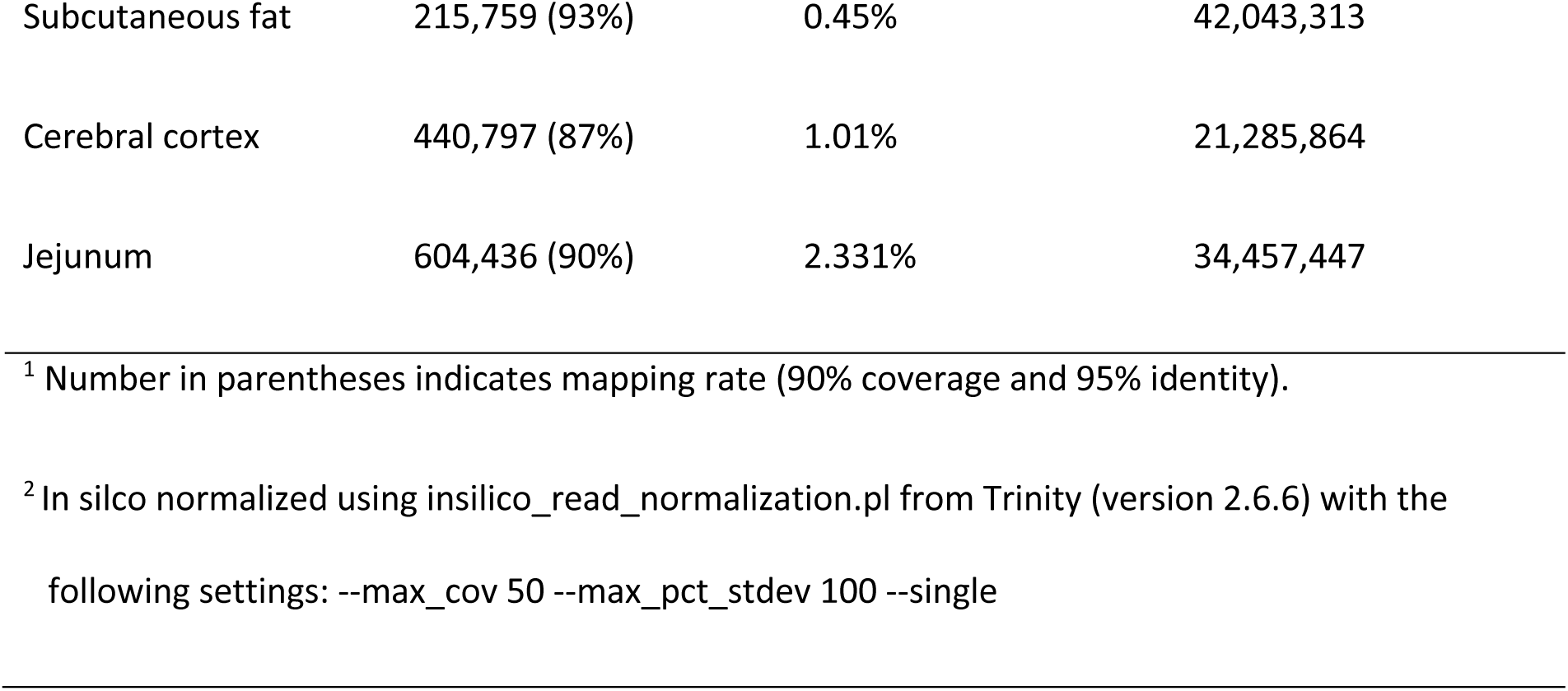
Summary of error-corrected, full-length non-chimeric (FLNC) Iso-Seq reads and their matched RNA-seq reads

| Tissue | Error-corrected FLNC<br>Iso-Seq reads<sup>1</sup> | Median error rate in<br>error-corrected FLNC<br>Iso-Seq reads | Normalized RNA-seq<br>reads used for error<br>correction<sup>2</sup> |
| --- | --- | --- | --- |
| Thalamus | 664,900 (90%) | 0.21% | 32,452,612 |
| Testes | 711,821 (86%) | 1.43% | 31,939,024 |
| Liver | 1,064,146 (84%) | 1.84% | 13,657,156 |
| Medulla | 380,531 (86%) | 0.43% | 48,256,918 |
| Subcutaneous fat | 215,759 (93%) | 0.45% | 42,043,313 |
| Cerebral cortex | 440,797 (87%) | 1.01% | 21,285,864 |
| Jejunum | 604,436 (90%) | 2.331% | 34,457,447 |

<sup>1</sup> Number in parentheses indicates mapping rate (90% coverage and 95% identity).
<sup>2</sup> In silico normalized using `insilico_read_normalization.pl` from Trinity (version 2.6.6) with the following settings: `--max_cov 50 --max_pct_stdev 100 --single`

### Prediction of transcript and gene biotypes

Transcripts’ open reading frames (ORFs) were predicted using the stand-alone version of ORFfinder (Wheeler et al. 2003) with “ATG and alternative initiation codons” as ORF start codon. The longest three ORFs were matched to the NCBI non-redundant vertebrate database and Uniprot vertebrate database using Blastp (Wheeler et al. 2003) with E-value cutoff of 10^− 6^, min coverage 60%, and min identity 95%. The ORFs with the lowest E-value to a protein were used as the representative, or if no matches were found, the longest ORF was used. Putative transcripts that had representative ORFs longer than 44 amino acids were labelled as protein-coding transcripts. If the representative ORF had a stop codon that was more than 50 bp upstream of the final splice junction, it was labelled as a nonsense-mediated decay transcript (Aken et al. 2016). Transcripts with start codon but no stop codon before their poly(A) site were labelled non-stop decay RNAs. Putative non-coding transcripts with lengths less than 200 bp were labelled as small non-coding RNAs (Aken et al. 2016). Putative non-coding transcripts with lengths greater than 200 bp were labelled as long non-coding RNAs (Aken et al. 2016). Long non-coding RNAs overlapping one or more coding loci on the opposite strand were labelled as antisense lncRNAs. Long non-coding RNAs located in introns of coding genes on the same strand were labelled as sense-intronic lncRNAs. Long non-coding RNAs that exonically overlapped with a protein-coding gene were labeled as Intragenic lncRNAs. Long non-coding RNAs located in intergenic regions of the genome were labeled as Intergenic lncRNAs.

Putative genes that transcribed at least a single protein-coding transcript were labelled as protein-coding genes. Putative genes with homology to existing vertebrate protein-coding genes (Blastx (Wheeler et al. 2003), E-value cut-off 10^-6^, min coverage 90%, and min identity 95%) but containing a disrupted coding sequence, i.e., transcribe only nonsense-mediated decay or non-stop decay transcripts in all of their detected tissues, were labelled as pseudogenes. The rest of the putative genes were labeled as non-coding.

Putative transcript structures were compared with independent bovine transcriptomes assembled from PacBio Iso-Seq data and RNA-seq data (see PacBio Iso-Seq data analysis), ONT-seq data (Halstead et al. 2021), and annotated transcripts from Ensembl (release 2021-03) and NCBI (Release 106) using Gffcompare (Pertea et al. 2016).

### ncRNAs homology analysis

Non-coding RNAs were matched to NCBI and Ensembl vertebrate ncRNA databases using Blastn (Wheeler et al. 2003) with E-value cutoff of 10^− 6^, min coverage 90%, and min identity 95%.

### Transcriptome termini site sequencing data analysis

T-rich stretches located at the 5^’^ end of each WTTS-seq raw read were removed using an in-house Perl script, as described previously (Zhou et al. 2016). T-trimmed reads were error-corrected using Coral (version 1.4.1) (Salmela and Schröder 2011) with -v -Y -u -a 3 option settings. The resulting reads were quality trimmed using FASTX Toolkit (version 0.0.14) (Hannon 2010) with -q 20 and -p 50 option settings. High-quality, error-corrected WTTS-seq reads were aligned against the ARS-UCD1.2 bovine genome using STAR (version 020201) (Dobin et al. 2013) with a cut-of of 95% identity and 90% coverage.

### ChIP-seq and ATAC-seq data analysis

The UC Davis FAANG Functional Annotation Pipeline was applied to process the ChIP-seq and ATAC-seq data, as previously described (Kern et al. 2021). Briefly, the ARS-UCD1.2 genome assembly and Ensembl genome annotation (v100) were used as references for cattle. Sequencing reads were trimmed with Trim Galore! (Krueger et al. 2015) (v.0.6.5) and aligned with either STAR (Dobin et al. 2012) (v.2.5.4a) or BWA (Li et al. 2013) (v0.7.17) to the respective genome assemblies. Alignments with MAPQ scores <30 were filtered using Samtools (Li et la. 2009) (v.1.9). For ChIP-seq, after the filtering, duplicates were marked and removed using Picard (v.2.18.7). Regions of signal enrichment (“peaks”) were called by MACS2 (Zhang et al. 2008) (v.2.1.1).

### Relating transcripts and genes to epigenetic data

The promoter was defined as the genomic region that spans from 500 bp 5ʹ to 100 bp 3ʹ of the gene/transcript start site. Histone mark or ATAC-seq (accesible chromatin) peaks mapped to the promoter of a given gene/transcript were related to that gene/transcript.

### Transcript and gene border validation

RAMPAGE peaks from a previously published experiment (Goszczynski et al. 2021) were used to validate gene/transcript start site. Peaks within the genomic region that spans from 30 bp 5ʹ to 10 bp 3ʹ of a gene/transcript start site were assigned to that gene/transcript. WTTS-seq reads (median length of 161 nt) within the genomic region that spans from 10 bp 5ʹ to 165 bp 3ʹ of a gene/transcript terminal site were assigned to that gene/transcript.

### Functional enrichment analysis

The potential mechanism of action of a group of genes was deciphered using ClueGO (Bindea et al. 2009). The latest update (May 2021) of the Gene Ontology Annotation database (GOA) (Huntley et al. 2015) was used in the analysis. The list of genes with at least one transcript detected in a given tissue was used as background for that tissue. The GO tree interval ranged from 3 to 20, with the minimum number of genes per cluster set to three. Term enrichment was tested with a right-sided hyper-geometric test that was corrected for multiple testing using the Benjamini-Hochberg procedure (Kim and van de Wiel 2008). The adjusted p-value threshold of 0.05 was used to filter enriched GO terms.

### Alternative splicing analysis

Alternative splicing events, except Unique Splice Site Exons, were detected using generateEvents from SUPPA (version 2.3) (Trincado et al. 2018) with default settings. Unique Splice Site Exons were detected using an in-house Python script.

### miRNA-seq data analysis

Single-end Qiagen miRNA-seq reads (50bp) from each tissue sample were trimmed to remove the adaptor sequences and low-quality bases using Trim Galore (version 0.6.4) (Krueger 2019) with --quality 20, --length 16, --max_length 30 -a AACTGTAGGCACCATCAAT option settings. miRNA reads were aligned against the ARS-UCD1.2 bovine genome using mapper.pl from mirDeep2 (version 0.1.3) (Friedländer et al. 2012) with -e -h -q -j -l 16 -o 40 -r 1 -m -v -n option settings. miRNA mature sequences along with their hairpin sequences for Bos taurus species were downloaded from miRbase (Kozomara et al. 2019). These sequences, along with the aligned miRNA reads, were used to quantify known miRNAs in each sample using miRDeep2.pl from mirDeep2 (version 0.1.3) (Friedländer et al. 2012) with -t bta -c -v 2 setting options. miRNA normalized Reads Per Million (RPM) were used to check sample similarities using hierarchical clustering and regression analysis of gene expression values (log2 based CPM), and outlier samples were detected and removed from downstream analysis. In order to predict the most comprehensive set of novel miRNAs, samples from different tissues were concatenated into a single file that were aligned against the ARS-UCD1.2 bovine genome using mapper.pl from mirDeep2 (version 0.1.3) (Friedländer et al. 2012) with the aforementioned settings. Aligned reads from the previous step were used, along with known miRNAs’ mature sequences and their hairpins, to predict novel miRNAs using miRDeep2.pl from mirDeep2 (version 0.1.3) (Friedländer et al. 2012) with the aforementioned settings. Samples from each tissue were combined to get the most comprehensive set of data for that tissue. Mature miRNA sequences and their hairpins for both known and predicted novel miRNAs’ sequences along with the aligned miRNA reads from each tissue were used to quantify known and novel miRNAs in each tissue using mirDeep2 (version 0.1.3) (Friedländer et al. 2012) with the aforementioned settings.

### Tissue-specificity index

Tissue Specificity Index (TSI) calculations were utilized to present more comprehensive information on transcript/gene/miRNA expression patterns across tissues. This index has a range of zero to one with a score of zero corresponding to ubiquitously expressed transcripts/genes/miRNAs (i.e., “housekeepers”) and a score of one for transcripts/genes/miRNAs that are expressed in a single tissue (i.e., “tissue-specific”) (Ludwig et al. 2016). The TSI for a transcript/gene/miRNA j was calculated as (Ludwig et al. 2016):

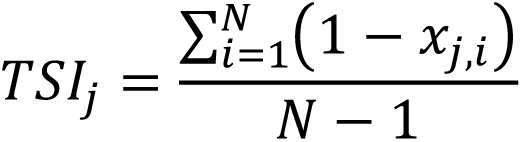

where *N* corresponds to the total number of tissues measured, and *x*_*j,i*_ is the expression intensity of tissue *i* normalized by the maximal expression of any tissue for transcript/gene/miRNA *j*.

### QTL enrichment analysis

Publicly available bovine QTLs were retrieved from AnimalQTLdb (Hu et al. 2019). Genes closest to a given trait’s QTLs were denoted as QTL-associated genes for that trait. The median distance of QTLs located outside gene borders to the closest gene was 51.9 kilobases and the maximum distance was 2.6 million bases. QTL enrichment was tested with a right-sided Fisher Exact test using an in-house Python script. The resulting p-values were corrected for multiple testing by the Benjamini-Hochberg procedure (Kim and van de Wiel 2008). The adjusted p-value threshold of 0.05 was used to filter QTLs.

### Trait similarity network

For a given pair of traits, trait A was denoted as “similar” to trait B if a significant portion of trait A’s QTL-associated genes were also the closest genes to trait B QTLs based on 1000 permutation tests. The resulting p-values were corrected for multiple testing using the Benjamini-Hochberg procedure (Kim and van de Wiel 2008). The same procedure was used to test trait B’s similarity to trait A. The adjusted p-value threshold of 0.05 was used to filter significant trait similarities. A graphical presentation of the method used to construct the tissue similarity network is presented in Supplemental file 1: Fig. S39. The resulting network was visualized using Cystoscape software (Shannon et al. 2003).

### Testis-pituitary axis correlation significance test

The presence of signal peptides on representative ORFs of protein-coding transcripts was predicted using SignalP-5.0 (Almagro Armenteros et al. 2019). Spearman correlation coefficients were used to study expression similarity between testis genes encoding signal peptides that were closest to the “percentage of normal sperm” QTLs (62 genes) and pituitary genes closest to the “percentage of normal sperm” QTLs (246 genes). To test the statistical difference between these correlation coefficients (reference correlations) and random chance, 1000 random sets of 246 pituitary genes were selected, and their correlation coefficients with 62 previously described testis genes were calculated (random correlations). The reference correlations were compared with 1000 sets of random correlations using a right-sided t-test.

The resulting p-values were corrected for multiple testing by the Benjamini-Hochberg procedure (Kim and van de Wiel 2008). The distribution-adjusted p-values were used to determine the significance level of expression similarities for genes involved in the testis-pituitary axis related to “percentage of normal sperm”. The same analysis was conducted to determine the significance of pituitary-testis axis involvement in this trait.

### Tissue dendrogram comparison across different transcript and gene biotypes

Tissues were clustered based on the percentage of their transcripts/genes that were shared between tissue pairs using the hclust function in R. Cophenetic distances for tissue dendrograms were calculated using the cophenetic R function. The degree of similarity between dendrograms constructed based on different gene/transcript biotypes was obtained using the Spearman correlation coefficient between the dendrograms’ Cophenetic distances.

## Supplemental files

**Supplemental file 1: Fig. S1** Distribution of the number of RNA-seq reads across tissues. **Fig. S2** (A) Comparison of tissues based on number of transcript biotypes and (B) percentage of transcript biotypes. (C) Comparison of transcript biotypes based on their number of detected tissues and (D) their expression level across detected tissues. **Fig. S3** (A) Relation between the number of input reads and the number of transcript biotypes (B) Comparison of expression level between different transcript biotypes. **Fig. S4** Tissue similarities (A) and clustering (B) based on the percentage of protein-coding transcripts shared between pairs of tissues. **Fig. S5** Tissue similarities (A) and clustering (B) based on the percentage of non-coding transcripts shared between pairs of tissues. **Fig. S6** Comparison of known and novel transcripts based on their expression (A) and number of detected tissues (B). **Fig. S7** Comparison of known and novel protein-coding transcripts based on the length (A) and GC content (B) of their 5’ UTR, CDS, and 3’ UTR. **Fig. S8** (A) Comparison of tissues based on number of gene biotypes and (B) percentage of gene biotypes. **Fig. S9** Relation between the number of input reads and the number of gene biotypes**. Fig. S10** Comparison of known and novel genes based on their expression (A) and number of detected tissues (B). **Fig. S11** Functional enrichment analysis of the top five percent of genes with the highest number of UTRs. **Fig. S12** Similarity of tissues based on the number of non-coding genes in their fetal samples that switched to protein-coding genes with only coding transcripts in their adult samples. **Fig. S13** (A) Distribution of genes that transcribed PATs, based on their number of detected tissues, percentage of genes’ transcripts that are PATs and percentage of genes’ detected tissues in which PATs were transcribed. (B) Comparison of genes that transcribed PATs with other gene biotypes. **Fig. S14** (A) Homology analysis of protein-coding genes. (B) Homology analysis of non-coding genes. (C) Detection of orphan genes based on homology classification of cattle-specific protein-coding genes and non-coding genes. **Fig. S15** Comparison of the expression level of homologous and orphan genes across (A) and within (B) their detected tissues. (C) Comparison of homologous and orphan genes based on the number of detected tissues. **Fig. S16** Comparison of different gene biotypes based on the expression (A) and the number of detected tissues (B). **Fig. S17** Comparison of different pseudogene-derived lncRNAs and non-pseudogene derived lncRNAs based on the expression level (A) and the number of detected tissues (B)**. Fig. S18** Tissue similarities (A) and clustering (B) based on the percentage of protein-coding genes shared between pairs of tissues. **Fig. S19** Tissue similarities (A) and clustering (B) based on the percentage of non-coding genes shared between pairs of tissues. **Fig. S20** (A) Different types of alternative splicing events. (B) Comparison of bovine genome builds based on the number of transcripts that showed any type of alternative splicing (AS) events**. Fig. S21** Comparison of tissues based on the number (A) and the percentage (B) of transcripts that showed different types of alternative splicing events. Comparison of tissues based on the number (C) and the percentage (D) of alternative splicing events**. Fig. S22** (A) Comparison of tissues based on the percentage of transcripts that showed any type of alternative splicing events, spliced transcripts from single-transcript genes, and unspliced transcripts and (B) the relation between the number of input reads and the number of these transcripts across tissues. **Fig. S23** Comparison of transcripts that showed different types of alternative splicing events based on (A) the expression level in the detected tissues and (B) the number of detected tissues. **Fig. S24** Transcript biotype switching due to alternative splicing events**. Fig. S25** Comparison of tissues based on the number of alternative splicing events per alternatively spliced gene**. Fig. S26** (A) Distribution of the number of alternative splicing events per alternatively spliced gene. The 5% quantile is shown using a dashed red line. (B) Functional enrichment analysis of the top five percent of genes with the highest number of alternative splicing events. **Fig. S27** Comparison of the alternative splicing rate between adult and fetal tissues. **Fig. S28** (A) Distribution of gene’s number of detected tissues. Tissue-specific gene biotypes are shown in the pie chart. (B) Distribution of transcript’s number of detected tissues. Tissue-specific transcript biotypes are shown in the pie chart. (C) Comparison of tissues based on the number of tissue-specific genes and transcripts. (D) Comparison of the expression level of tissue-specific genes and transcripts versus their non-tissue-specific counterparts. **Fig. S29** Relationship between tissue specificity and alternative splicing events**. Fig. S30** Relationship between tissue specificity index and the number of multi-tissue detected genes (A) and transcripts (B). Distribution of tissue specificity indexes in multi-tissue detected genes (C) and transcripts (D). The 5% quantile is shown using dashed red lines. (E) Functional enrichment analysis of the top five percent of multi-tissue detected genes with the highest tissue specificity indexes. **Fig. S31** Distribution of QTLs located outside gene borders in relation to the closest gene**. Fig. S32** (A) Distribution of correlation coefficients between SPACA5 gene expression and pituitary genes closest to “percentage of normal sperm” QTLs. Dashed lines show the minimum significant positive and negative correlation (p-value <0.05). (B) Expression atlas of SPACA5 gene in human tissues from The Human Protein Atlas (Uhlén et al. 2015). **Fig. S33** (A) Correlation between pituitary genes with signal peptides that were close to the “percentage of normal sperm” QTL and testis genes closest to this trait’s QTL (reference correlations). (B) Distribution of p-values resulting from right-sided t-test between reference correlation coefficients and correlation coefficients derived from random chance (see methods for details)**. Fig. S34** Tissue similarities (A) and clustering (B) based on the percentage of miRNAs shared between pairs of tissues. **Fig. S35** Clustering of tissues based on protein-coding genes (A), protein-coding transcripts (B), non-coding genes (C), non-coding transcripts (D), and miRNAs (E). (F) Comparison of tissue dendrograms based on the correlation between their Cophenetic distances. **Fig. S36** (A) Distribution of the number of detected tissues for known and novel miRNAs. Classification of miRNAs as known, or novel is presented in the pie chart. (B) Comparison of tissues based on their number of tissue-specific miRNAs. (C) Expression of known and novel miRNAs in their detected tissues. (D) Distribution of multi-tissue detected miRNAs’ tissue specificity indexes. (E) Relationship between tissue specificity index and number of detected tissues in multi-tissue detected miRNAs. Dots have been color coded based on their density. **Fig. S37** Distribution of the number of detected genes (A), transcripts (B), and miRNAs (C) across tissues. **Fig. S38** Distribution of the number of known and novel genes (A), transcripts (B), and miRNAs (C) across tissues. **Fig. S39** Graphical representation of the method used to construct the tissue similarity network.

**Supplemental file 2:** Summary of RNA-seq and miRNA-seq reads

**Supplemental file 3:** Detailed description of the number of transcripts, genes, and miRNAs detected in each tissue

**Supplemental file 4:** List of transcripts and genes expressed in each tissue and their expression values (RPKM)

**Supplemental file 5:** Transcript biotype enrichment analysis in adult and fetal tissues

**Supplemental file 6:** Functional enrichment analysis of the top five percent of genes with the highest number of UTRs

**Additional file7:** Functional enrichment analysis of genes that remained bifunctional in all their detected tissues

**Additional file8:** Functional enrichment analysis of non-coding genes in fetal tissues that were switched to protein coding with only coding transcripts in their matched adult tissue

**Additional file9:** Functional enrichment analysis of protein-coding genes that transcribed PATs as their main transcripts (PATs comprised >50% of their transcripts) in all their detected tissues

**Supplemental file 10:** Gene biotype enrichment analysis in adult and fetal tissues

**Supplemental file 11:** Functional enrichment analysis of the top five percent of genes with the highest number of alternative splicing events

**Additional file12:** List of tissue-specific genes and transcripts

**Additional file13:** Genes and transcripts tissue specificity indexes

**Additional file14:** Functional enrichment analysis of the top five percent of multi-tissue detected genes with the highest tissue specificity indexes

**Additional file15:** List of QTL’s closest genes in each tissue

**Additional file16:** Trait enrichment analysis of testis-specific genes

**Additional file16:** Pituitary genes closest to “percentage of normal sperm” QTLs that showed positive significant correlation with SPACA5 gene in testis

**Additional file18:** List of genes closest to “percentage of normal sperm” QTLs that were involved in testis-pituitary tissue axis and their functional enrichment analysis results

**Additional file19:** List of genes closest to “percentage of normal sperm” QTLs that were involved in pituitary-testis tissue axis and their functional enrichment analysis results

**Additional file20:** Similarity of traits based on the integration of the assembled bovine transcriptome with publicly available QTLs

**Additional file21:** List of miRNAs expressed in each tissue and their expression values

**Additional file22:** List of tissues related to different omics datasets used in the experiment

## Data availability

RNA-seq and miRNA-seq, ATAC-seq, and WTTS-seq datasets generated in this study are submitted to the ArrayExpress database (https://www.ebi.ac.uk/arrayexpress/) under accession numbers MTAB-11699, E-MTAB-11815, and E-MTAB-12052, respectively. PacBio Iso-Seq, ChIP-seq, and RAMPAGE datasets generated in this study are submitted to the NCBI BioProject database (https://www.ncbi.nlm.nih.gov/bioproject/) under accession numbers PRJNA386670, GSE158416, and PRJNA630504, respectively. The constructed bovine trait similarity network is publicly available through the Animal Genome database (https://www.animalgenome.org/host/reecylab/a). The constructed cattle transcriptome and related sequences are publicly available in the Open Science Framework database (https://osf.io/jze72/?view_only=d2dd1badf37e4bafae1e12731a0cc40d). Custom code used is available at https://github.com/hamidbeiki/Cattle-Genome.

## Ethics approval and consent to participate

Procedures for tissue collection followed the Animal Care and Use protocol (#18464) approved by the Institutional Animal Care and Use Committee (IACUC), University of California, Davis (UCD).

## Consent for publication

Not applicable

## Competing interests

The authors declare no competing interests.

## Funding

This study was supported by Agriculture and Food Research Initiative Competitive Grant no. 2018-67015-27500 (H.Z., P.R. etc.) and sample collection was supported by no. 2015-67015-22940 (H.Z. and P.R.) from the USDA National Institute of Food and Agriculture.

## Acknowledgments

We are grateful to Nathan Weeks for helping with massive parallel computing of transcriptome assembly.

## Authors’ contributions

H.B., B.M.M., H.J., H.Z., M.R., P.J.R., S.M., T.P.L.S., W.L., Z.J., and J.M.R. conceived and designed the project; C.K., W.M., and W.L. generated RNA-seq and miRNA-seq data; D.K., G.B., J.T., and K.D. participated in tissue collection; R.H and H.J prepared cells; J.J.M., X.Z., X.H., and Z.J. generated W.T.T.S-seq data, X.X., P.J.R. and H.J generated ChIP-seq data; M.R.J. generated ATAC-seq data; T.P.L.S. generated PacBio Iso-seq data; G.R. and S.C. conducted sequencing of RNA-seg, miRNA-seq, ChIP-seq, and ATAC-seq data; H.B. conducted bioinformatics data analysis and drafted the manuscript, which was edited by C.A.P., B.M.M., H.J., H.Z., J.E.K., M.R., P.J.R., S.M., T.P.L.S., W.L., Z.J. and J.M.R.; Z.H. created the web-based database for the trait similarity network; all authors read and approved the final manuscript.

## Endnotes

Mention of trade names or commercial products in this publication is solely for the purpose of providing specific information and does not imply recommendation or endorsement by the U.S. Department of Agriculture. USDA is an equal opportunity provider and employer.

The results reported here were made possible with resources provided by the USDA shared computing cluster (Ceres) as part of the ARS SCINet initiative.

## Notes

### Competing Interest Statement

The authors have declared no competing interest.

## References

Aken BL, Ayling S, Barrell D, Clarke L, Curwen V, Fairley S, Fernandez Banet J, Billis K, García Girón C, Hourlier T et al. 2016. The Ensembl gene annotation system. Database (Oxford) 2016.

Almagro Armenteros JJ, Tsirigos KD, Sønderby CK, Petersen TN, Winther O, Brunak S, von Heijne G, Nielsen H. 2019. SignalP 5.0 improves signal peptide predictions using deep neural networks. Nature Biotechnology 37: 420–423.

Ambros V. 2004. The functions of animal microRNAs. Nature 431: 350–355.

Anderson DM, Anderson KM, Chang CL, Makarewich CA, Nelson BR, McAnally JR, Kasaragod P, Shelton JM, Liou J, Bassel-Duby R et al. 2015. A micropeptide encoded by a putative long noncoding RNA regulates muscle performance. Cell 160: 595–606.

Andrews SJ, Rothnagel JA. 2014. Emerging evidence for functional peptides encoded by short open reading frames. Nat Rev Genet 15: 193–204.

Araujo PR, Yoon K, Ko D, Smith AD, Qiao M, Suresh U, Burns SC, Penalva LO. 2012. Before It Gets Started: Regulating Translation at the 5’ UTR. Comp Funct Genomics 2012: 475731.

Bartel DP. 2004. MicroRNAs: genomics, biogenesis, mechanism, and function. Cell 116: 281–297.

Beiki H, Liu H, Huang J, Manchanda N, Nonneman D, Smith TPL, Reecy JM, Tuggle CK. 2019. Improved annotation of the domestic pig genome through integration of Iso-Seq and RNA-seq data. BMC Genomics 20: 344.

Bindea G, Mlecnik B, Hackl H, Charoentong P, Tosolini M, Kirilovsky A, Fridman WH, Pagès F, Trajanoski Z, Galon J. 2009. ClueGO: a Cytoscape plug-in to decipher functionally grouped gene ontology and pathway annotation networks. Bioinformatics 25: 1091–1093.

Chen J, Chen ZJ. 2013. Regulation of NF-κB by ubiquitination. Curr Opin Immunol 25: 4–12.

Clare M, Richard P, Kate K, Sinead W, Mark M, David K. 2018. Residual feed intake phenotype and gender affect the expression of key genes of the lipogenesis pathway in subcutaneous adipose tissue of beef cattle. J Anim Sci Biotechnol 9: 68.

Colombo M, Karousis ED, Bourquin J, Bruggmann R, Mühlemann O. 2017. Transcriptome-wide identification of NMD-targeted human mRNAs reveals extensive redundancy between SMG6- and SMG7-mediated degradation pathways. RNA 23: 189–201.

Dobin A, Davis CA, Schlesinger F, Drenkow J, Zaleski C, Jha S, Batut P, Chaisson M, Gingeras TR. 2013. STAR: ultrafast universal RNA-seq aligner. Bioinformatics 29: 15–21.

Elolimy AA, Abdelmegeid MK, McCann JC, Shike DW, Loor JJ. 2018. Residual feed intake in beef cattle and its association with carcass traits, ruminal solid-fraction bacteria, and epithelium gene expression. J Anim Sci Biotechnol 9: 67.

Farr VC, Stelwagen K, Cate LR, Molenaar AJ, McFadden TB, Davis SR. 1996. An improved method for the routine biopsy of bovine mammary tissue. J Dairy Sci 79: 543–549.

Friedländer MR, Mackowiak SD, Li N, Chen W, Rajewsky N. 2012. miRDeep2 accurately identifies known and hundreds of novel microRNA genes in seven animal clades. Nucleic Acids Res 40: 37–52.

Gerber S, Schratt G, Germain PL. 2021. Streamlining differential exon and 3’ UTR usage with diffUTR. BMC Bioinformatics 22: 189.

Gonzàlez-Porta M, Frankish A, Rung J, Harrow J, Brazma A. 2013. Transcriptome analysis of human tissues and cell lines reveals one dominant transcript per gene. Genome Biol 14: R70.

Goszczynski DE, Halstead MM, Islas-Trejo AD, Zhou H, Ross PJ. 2021. Transcription initiation mapping in 31 bovine tissues reveals complex promoter activity, pervasive transcription, and tissue-specific promoter usage. Genome Res 31: 732–744.

Grabherr MG, Haas BJ, Yassour M, Levin JZ, Thompson DA, Amit I, Adiconis X, Fan L, Raychowdhury R, Zeng Q et al. 2011. Full-length transcriptome assembly from RNA-Seq data without a reference genome. Nat Biotechnol 29: 644–652.

Green TC, Jago JG, Macdonald KA, Waghorn GC. 2013. Relationships between residual feed intake, average daily gain, and feeding behavior in growing dairy heifers. J Dairy Sci 96: 3098–3107.

Hackl T, Hedrich R, Schultz J, Förster F. 2014. proovread: large-scale high-accuracy PacBio correction through iterative short read consensus. Bioinformatics 30: 3004–3011.

Halasa T, Kirkeby C. 2020. Differential Somatic Cell Count: Value for Udder Health Management. Front Vet Sci 7: 609055.

Halstead MM, Islas-Trejo A, Goszczynski DE, Medrano JF, Zhou H, Ross PJ. 2021. Large-Scale Multiplexing Permits Full-Length Transcriptome Annotation of 32 Bovine Tissues From a Single Nanopore Flow Cell. Front Genet 12: 664260.

Hannon GJ. 2010. FASTX-Toolkit. http://hannonlab.cshl.edu/fastx_toolkit.

Hass B. 2015. https://hpcgridrunner.github.io/.

Hertl JA, Schukken YH, Tauer LW, Welcome FL, Gröhn YT. 2018. Does clinical mastitis in the first 100 days of lactation 1 predict increased mastitis occurrence and shorter herd life in dairy cows? J Dairy Sci 101: 2309–2323.

Hindorff LA, Sethupathy P, Junkins HA, Ramos EM, Mehta JP, Collins FS, Manolio TA. 2009. Potential etiologic and functional implications of genome-wide association loci for human diseases and traits. Proc Natl Acad Sci U S A 106: 9362–9367.

Houlahan K, Schenkel FS, Hailemariam D, Lassen J, Kargo M, Cole JB, Connor EE, Wegmann S, Junior O, Miglior F et al. 2021. Effects of Incorporating Dry Matter Intake and Residual Feed Intake into a Selection Index for Dairy Cattle Using Deterministic Modeling. Animals (Basel) 11.

Hu ZL, Park CA, Reecy JM. 2019. Building a livestock genetic and genomic information knowledgebase through integrative developments of Animal QTLdb and CorrDB. Nucleic Acids Res 47: D701–D710.

Hubé F, Velasco G, Rollin J, Furling D, Francastel C. 2011. Steroid receptor RNA activator protein binds to and counteracts SRA RNA-mediated activation of MyoD and muscle differentiation. Nucleic Acids Res 39: 513–525.

Huntley RP, Sawford T, Mutowo-Meullenet P, Shypitsyna A, Bonilla C, Martin MJ, O’Donovan C. 2015. The GOA database: gene Ontology annotation updates for 2015. Nucleic Acids Res 43: D1057–1063.

Jereb S, Hwang HW, Van Otterloo E, Govek EE, Fak JJ, Yuan Y, Hatten ME, Darnell RB. 2018. Differential 3’ Processing of Specific Transcripts Expands Regulatory and Protein Diversity Across Neuronal Cell Types. Elife 7.

Kaniyamattam K, De Vries A, Tauer LW, Gröhn YT. 2020. Economics of reducing antibiotic usage for clinical mastitis and metritis through genomic selection. J Dairy Sci 103: 473–491.

Karalis KP, Venihaki M, Zhao J, van Vlerken LE, Chandras C. 2004. NF-kappaB participates in the corticotropin-releasing, hormone-induced regulation of the pituitary proopiomelanocortin gene. J Biol Chem 279: 10837–10840.

Kern C, Wang Y, Xu X, Pan Z, Halstead M, Chanthavixay G, Saelao P, Waters S, Xiang R, Chamberlain A et al. 2021. Functional annotations of three domestic animal genomes provide vital resources for comparative and agricultural research. Nat Commun 12: 1821.

Kim KI, van de Wiel MA. 2008. Effects of dependence in high-dimensional multiple testing problems. BMC Bioinformatics 9: 114.

Kozomara A, Birgaoanu M, Griffiths-Jones S. 2019. miRBase: from microRNA sequences to function. Nucleic Acids Res 47: D155–D162.

Krueger F. 2019. https://www.bioinformatics.babraham.ac.uk/projects/trim_galore/.

Kumar S, Lee HJ, Park HS, Lee K. 2016. Testis-Specific GTPase (TSG): An oligomeric protein. BMC Genomics 17: 792.

Kumari P, Sampath K. 2015. cncRNAs: Bi-functional RNAs with protein coding and non-coding functions. Semin Cell Dev Biol 47-48: 40-51.

Kurosaki T, Popp MW, Maquat LE. 2019. Quality and quantity control of gene expression by nonsense-mediated mRNA decay. Nat Rev Mol Cell Biol 20: 406–420.

Leek J, Johnson W, Parker HS, Fertig EJ, Jaffe AE, Zhang Y, Storey JD, LC T. 2021. sva: Surrogate Variable Analysis. R package version 3.30.0.

Li J, Liu C. 2019. Coding or Noncoding, the Converging Concepts of RNAs. Front Genet 10: 496.

Liao Y, Smyth GK, Shi W. 2014. featureCounts: an efficient general purpose program for assigning sequence reads to genomic features. Bioinformatics 30: 923–930.

Lima FS, Silvestre FT, Peñagaricano F, Thatcher WW. 2020. Early genomic prediction of daughter pregnancy rate is associated with improved reproductive performance in Holstein dairy cows. J Dairy Sci 103: 3312–3324.

Liu E, VandeHaar MJ. 2020. Relationship of residual feed intake and protein efficiency in lactating cows fed high- or low-protein diets. J Dairy Sci 103: 3177–3190.

Lou W, Ding B, Fu P. 2020. Pseudogene-Derived lncRNAs and Their miRNA Sponging Mechanism in Human Cancer. Front Cell Dev Biol 8: 85.

Ludwig N, Leidinger P, Becker K, Backes C, Fehlmann T, Pallasch C, Rheinheimer S, Meder B, Stähler C, Meese E et al. 2016. Distribution of miRNA expression across human tissues. Nucleic Acids Res 44: 3865–3877.

Mackowiak SD, Zauber H, Bielow C, Thiel D, Kutz K, Calviello L, Mastrobuoni G, Rajewsky N, Kempa S, Selbach M et al. 2015. Extensive identification and analysis of conserved small ORFs in animals. Genome Biol 16: 179.

Martí De Olives A, Díaz JR, Molina MP, Peris C. 2013. Quantification of milk yield and composition changes as affected by subclinical mastitis during the current lactation in sheep. J Dairy Sci 96: 7698–7708.

Mayba O, Gilbert HN, Liu J, Haverty PM, Jhunjhunwala S, Jiang Z, Watanabe C, Zhang Z. 2014. MBASED: allele-specific expression detection in cancer tissues and cell lines. Genome Biol 15: 405.

Mazin PV, Khaitovich P, Cardoso-Moreira M, Kaessmann H. 2021. Alternative splicing during mammalian organ development. Nature Genetics 53: 925–934.

Miles AM, McArt JAA, Leal Yepes FA, Stambuk CR, Virkler PD, Huson HJ. 2019. Udder and teat conformational risk factors for elevated somatic cell count and clinical mastitis in New York Holsteins. Prev Vet Med 163: 7–13.

Milligan MJ, Lipovich L. 2014. Pseudogene-derived lncRNAs: emerging regulators of gene expression. Front Genet 5: 476.

Mitrovich QM, Anderson P. 2005. mRNA surveillance of expressed pseudogenes in C. elegans. Curr Biol 15: 963–967.

Nam JW, Choi SW, You BH. 2016. Incredible RNA: Dual Functions of Coding and Noncoding. Mol Cells 39: 367–374.

Nickless A, Bailis JM, You Z. 2017. Control of gene expression through the nonsense-mediated RNA decay pathway. Cell Biosci 7: 26.

O’Shaughnessy PJ, Fleming LM, Jackson G, Hochgeschwender U, Reed P, Baker PJ. 2003. Adrenocorticotropic hormone directly stimulates testosterone production by the fetal and neonatal mouse testis. Endocrinology 144: 3279–3284.

Olexiouk V, Crappé J, Verbruggen S, Verhegen K, Martens L, Menschaert G. 2016. sORFs.org: a repository of small ORFs identified by ribosome profiling. Nucleic Acids Res 44: D324–329.

PacificBiosciences. 2018. https://www.pacb.com/products-and-services/analytical-software/smrt-analysis/.

Pedersen BS, Quinlan AR. 2018. Mosdepth: quick coverage calculation for genomes and exomes. Bioinformatics 34: 867–868.

Pertea M, Kim D, Pertea GM, Leek JT, Salzberg SL. 2016. Transcript-level expression analysis of RNA-seq experiments with HISAT, StringTie and Ballgown. Nat Protoc 11: 1650–1667.

Rajala-Schultz PJ, Gröhn YT, McCulloch CE, Guard CL. 1999. Effects of clinical mastitis on milk yield in dairy cows. J Dairy Sci 82: 1213–1220.

Remnant J, Green MJ, Huxley J, Hirst-Beecham J, Jones R, Roberts G, Hudson CD. 2019. Association of lameness and mastitis with return-to-service oestrus detection in the dairy cow. Vet Rec 185: 442.

Richburg JH, Myers JL, Bratton SB. 2014. The role of E3 ligases in the ubiquitin-dependent regulation of spermatogenesis. Semin Cell Dev Biol 30: 27–35.

Roth JA, Tuggle CK. 2015. Livestock models in translational medicine. ILAR J 56: 1–6.

Salmela L, Schröder J. 2011. Correcting errors in short reads by multiple alignments. Bioinformatics 27: 1455–1461.

Sammeth M, Foissac S, Guigó R. 2008. A general definition and nomenclature for alternative splicing events. PLoS Comput Biol 4: e1000147.

Schurch NJ, Cole C, Sherstnev A, Song J, Duc C, Storey KG, McLean WH, Brown SJ, Simpson GG, Barton GJ. 2014. Improved annotation of 3’ untranslated regions and complex loci by combination of strand-specific direct RNA sequencing, RNA-Seq and ESTs. PLoS One 9: e94270.

Shannon P, Markiel A, Ozier O, Baliga NS, Wang JT, Ramage D, Amin N, Schwikowski B, Ideker T. 2003. Cytoscape: a software environment for integrated models of biomolecular interaction networks. Genome Res 13: 2498–2504.

Stewart GL, Enfield KSS, Sage AP, Martinez VD, Minatel BC, Pewarchuk ME, Marshall EA, Lam WL. 2019. Aberrant Expression of Pseudogene-Derived lncRNAs as an Alternative Mechanism of Cancer Gene Regulation in Lung Adenocarcinoma. Front Genet 10: 138.

Supek F, Lehner B, Lindeboom RGH. 2021. To NMD or Not To NMD: Nonsense-Mediated mRNA Decay in Cancer and Other Genetic Diseases. Trends Genet 37: 657–668.

Tange O. 2018. GNU Parallel. https://doi.org/10.5281/zenodo.1146014.

Tixier-Boichard M, Fabre S, Dhorne-Pollet S, Goubil A, Acloque H, Vincent-Naulleau S, Ross P, Wang Y, Chanthavixay G, Cheng H et al. 2021. Tissue Resources for the Functional Annotation of Animal Genomes. Front Genet 12: 666265.

Trincado JL, Entizne JC, Hysenaj G, Singh B, Skalic M, Elliott DJ, Eyras E. 2018. SUPPA2: fast, accurate, and uncertainty-aware differential splicing analysis across multiple conditions. Genome Biol 19: 40.

Uhlén M, Fagerberg L, Hallström BM, Lindskog C, Oksvold P, Mardinoglu A, Sivertsson Å, Kampf C, Sjöstedt E, Asplund A et al. 2015. Proteomics. Tissue-based map of the human proteome. Science 347: 1260419.

Wang JR, Holt J, McMillan L, Jones CD. 2018. FMLRC: Hybrid long read error correction using an FM-index. BMC Bioinformatics 19: 50.

Weber C, Hametner C, Tuchscherer A, Losand B, Kanitz E, Otten W, Singh SP, Bruckmaier RM, Becker F, Kanitz W et al. 2013. Variation in fat mobilization during early lactation differently affects feed intake, body condition, and lipid and glucose metabolism in high-yielding dairy cows. J Dairy Sci 96: 165–180.

Wei L-H, Guo JU. 2020. Coding functions of “noncoding” RNAs. Science 367: 1074–1075.

Wheeler DL, Church DM, Federhen S, Lash AE, Madden TL, Pontius JU, Schuler GD, Schriml LM, Sequeira E, Tatusova TA et al. 2003. Database resources of the National Center for Biotechnology. Nucleic Acids Res 31: 28–33.

Wollerton MC, Gooding C, Wagner EJ, Garcia-Blanco MA, Smith CW. 2004. Autoregulation of polypyrimidine tract binding protein by alternative splicing leading to nonsense-mediated decay. Mol Cell 13: 91–100.

Yates LA, Norbury CJ, Gilbert RJ. 2013. The long and short of microRNA. Cell 153: 516–519.

Yi Z, Li X, Luo W, Xu Z, Ji C, Zhang Y, Nie Q, Zhang D, Zhang X. 2018. Feed conversion ratio, residual feed intake and cholecystokinin type A receptor gene polymorphisms are associated with feed intake and average daily gain in a Chinese local chicken population. J Anim Sci Biotechnol 9: 50.

Zhou X, Li R, Michal JJ, Wu XL, Liu Z, Zhao H, Xia Y, Du W, Wildung MR, Pouchnik DJ et al. 2016. Accurate Profiling of Gene Expression and Alternative Polyadenylation with Whole Transcriptome Termini Site Sequencing (WTTS-Seq). Genetics 203: 683–697.

